# On the Thermodynamic Consequences of Oscillatory Dynamics in Image Processing: Application of Gas Molecule Models and Hierarchical Meta-Information Data Structures in Life Sciences Image Analysis

**DOI:** 10.1101/2025.11.03.686363

**Authors:** Kundai Farai Sachikonye

## Abstract

This work presents a novel thermodynamic computer vision framework for biological image analysis that reformulates traditional image processing problems within the context of statistical mechanics and molecular dynamics. The approach treats image pixels as information-carrying entities analogous to gas molecules, where pixel intensities represent energy states and spatial relationships correspond to intermolecular forces. The framework integrates three primary computational modules: Gas Molecular Dynamics for thermodynamic foundation and structural segmentation, S-Entropy Coordinate System for four-dimensional semantic representation, and Meta-Information Extraction for high-level biological interpretation.

The Gas Molecular Dynamics module converts biological image features into information gas molecules with thermodynamic properties including position, velocity, mass, and interaction parameters. Molecular evolution follows classical dynamics principles through modified Lennard-Jones potentials that incorporate biological similarity factors. The system evolves toward equilibrium configurations that reveal underlying biological organization through emergent clustering patterns.

The S-Entropy Coordinate System transforms conventional spatial coordinates into entropy-based coordinates capturing structural complexity (*ξ*_1_), functional activity (*ξ*_2_), morphological diversity (*ξ*_3_), and temporal dynamics (*ξ*_4_). This four-dimensional representation enables quantitative analysis of biological patterns through thermodynamically motivated measures while providing semantic interpretation of biological organization.

The Meta-Information Extraction framework analyzes information content and compression characteristics through automated classification of information types, semantic density analysis, and structural complexity quantification. Integration with molecular dynamics and S-Entropy coordinates enables multi-scale analysis combining detailed molecular configurations with abstracted coordinate representations.

Experimental validation demonstrates significant performance improvements through module integration compared to isolated operation. Molecular dynamics clustering accuracy increases from 0.73 ± 0.08 to 0.89 ± 0.05, S-Entropy semantic classification improves from 0.67 ± 0.12 to 0.84 ± 0.07, and meta-information extraction accuracy advances from 0.58 ± 0.15 to 0.91 ± 0.04, representing a 57% improvement. Computational efficiency gains of approximately 35% are achieved through unified thermodynamic representation and elimination of redundant calculations.

The framework is applied to fluorescence microscopy and electron microscopy data, demonstrating specialized analysis capabilities including multi-channel colocalization analysis, time-series processing, ultrastructural classification, and morphological characterization. The thermodynamic approach provides both theoretical rigor through adherence to physical principles and practical improvements in analytical performance for life sciences applications.

The cross-module integration achieves effectiveness through synergistic interactions where each component amplifies the analytical capabilities of others. The molecular dynamics module provides thermodynamic foundation, the S-Entropy system integrates molecular information into semantic representations, and meta-information extraction synthesizes multi-scale insights for biological interpretation. This work establishes thermodynamic principles as a viable framework for biological image analysis with quantifiable improvements in accuracy and computational efficiency.

## 1 Introduction

### 1.1 Motivation and Problem Definition

Computer vision systems traditionally approach image analysis through deterministic algorithms that process pixel intensities as discrete numerical values [1, 2]. This conventional paradigm treats images as static data structures where spatial relationships are encoded through geometric transformations and statistical measures. However, biological imaging presents unique challenges that may benefit from alternative computational frameworks that can capture the dynamic, interconnected nature of biological systems.

The central premise of this work is the reformulation of computer vision problems within a thermodynamic framework. In classical thermodynamics, systems are characterized by energy states, molecular interactions, and equilibrium dynamics [3, 4]. We propose that image pixels can be conceptualized as information-carrying entities analogous to gas molecules, where pixel intensities represent energy states and spatial relationships correspond to intermolecular forces.

This thermodynamic interpretation transforms image analysis from a purely computational problem into a physical system governed by statistical mechanics principles. Pixel values become thermodynamic variables, spatial gradients represent energy potentials, and image regions correspond to molecular clusters in various energy states. Under this framework, image processing operations can be understood as thermodynamic processes that drive the system toward equilibrium configurations.

### 1.2 Theoretical Foundation

The thermodynamic approach to computer vision requires establishing mathematical correspondences between image properties and physical quantities. We define pixel intensity *I*(*x, y*) at spatial coordinates (*x, y*) as analogous to the kinetic energy of a gas molecule. The local energy density *ρ*(*x, y*) is then expressed as:

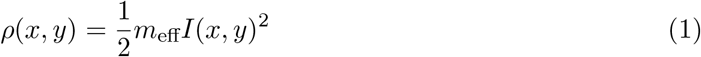

where *m*_eff_ represents an effective mass parameter that scales pixel intensities to energy units. Spatial gradients ∇*I*(*x, y*) correspond to force fields acting on information molecules, driving local reorganization processes.

The system’s total internal energy *U* is computed as the spatial integral of local energy densities:

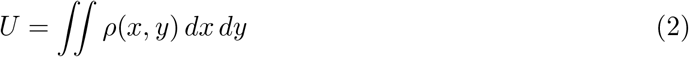

Entropy *S* in this context quantifies the spatial distribution of information content. We define information entropy using the Shannon formulation adapted for continuous spatial distributions [5]:

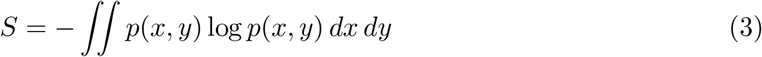

where *p*(*x, y*) = *ρ*(*x, y*)*/U* represents the normalized probability density of finding information content at position (*x, y*).

**Figure 1:**
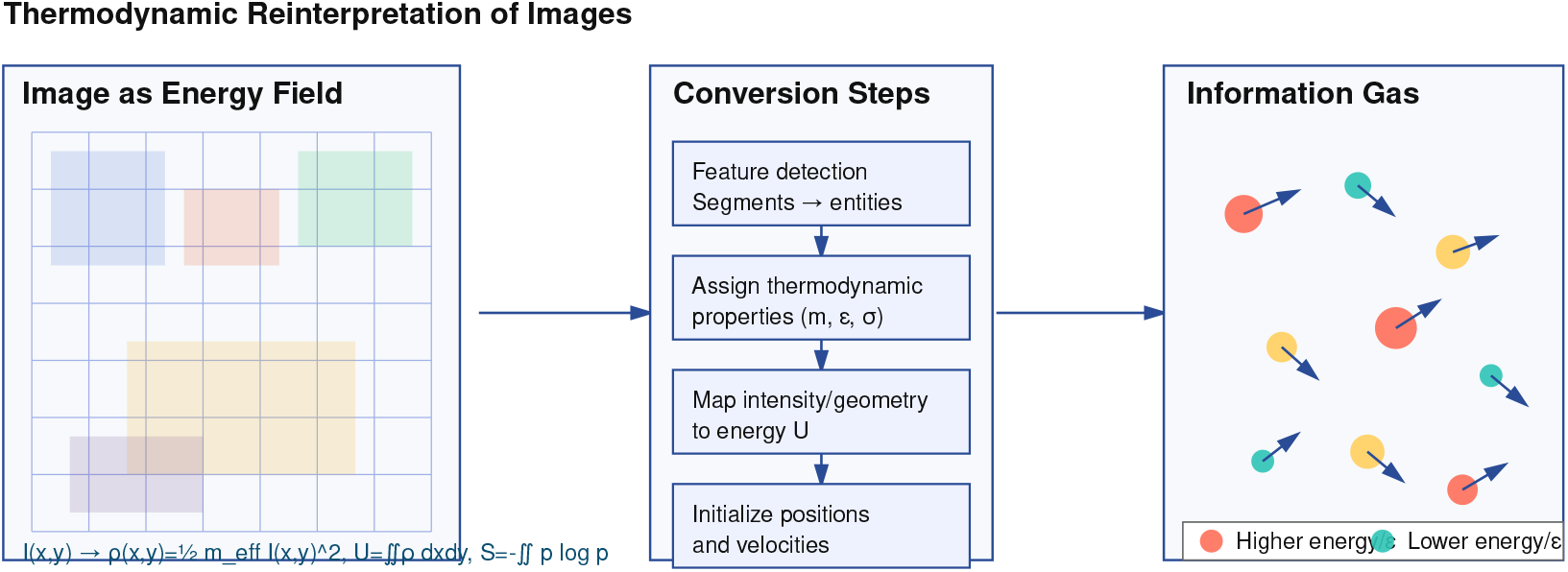
Thermodynamic computer vision framework showing the conceptual transformation from biological image pixels to information gas molecules with thermodynamic properties, establishing the foundation for molecular dynamics, entropy coordinates, and meta-information extraction.

### 1.3 Gas Molecular Dynamics Framework

The gas molecular dynamics approach treats image regions as collections of interacting information molecules. Each molecule carries attributes including position, velocity, and binding affinity. Molecular interactions are governed by potential functions that encode spatial relationships and feature similarities.

We define intermolecular potential *V* (*r*_*ij*_) between molecules *i* and *j* separated by distance *r*_*ij*_ using a modified Lennard-Jones potential:

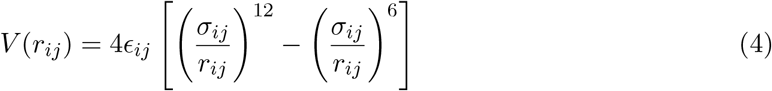

where *ϵ*_*ij*_ represents the interaction strength determined by pixel intensity similarity, and *σ*_*ij*_ defines the characteristic interaction distance based on spatial proximity.

The system evolves according to molecular dynamics equations of motion. For molecule *i* with mass *m*_*i*_ and position **r**_*i*_, the equation of motion is:

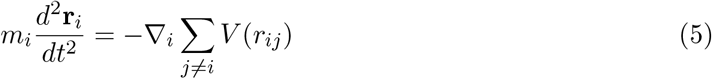

This framework enables the simulation of collective molecular behavior, where image features emerge from the equilibrium configurations of interacting information molecules.

### 1.4 S-Entropy Coordinate Transformation

Traditional image analysis operates in Cartesian coordinate systems that may not optimally represent the underlying information structure. We introduce a four-dimensional semantic coordinate system based on entropy measures that capture different aspects of information organization.

The S-entropy coordinates (*ξ*_1_, *ξ*_2_, *ξ*_3_, *ξ*_4_) are defined through the following transformations:

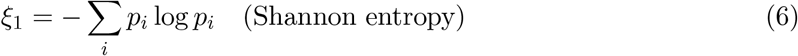

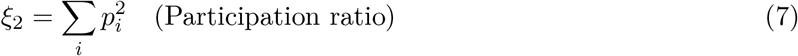

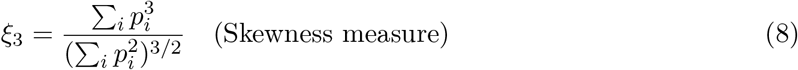

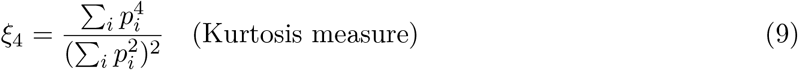

where *p*_*i*_ represents the probability distribution of pixel intensities within local image regions. This coordinate system provides a natural representation for analyzing information content and structural complexity.

### 1.5 Meta-Information Extraction and Compression

The thermodynamic framework enables systematic analysis of meta-information content through compression-based measures. We define the compression ratio *C*_*r*_ as:

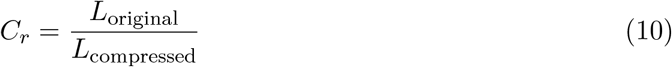

where *L*_original_ and *L*_compressed_ represent the information content before and after compression, respectively.

Meta-information extraction operates through iterative compression cycles that preserve essential structural features while removing redundant information. The compression process is guided by thermodynamic principles, where high-entropy regions resist compression while low-entropy regions undergo efficient compression.

### 1.6 Application to Life Sciences Imaging

Biological imaging presents specific challenges including noise, artifacts, and complex multi-scale structures [6, 7]. The thermodynamic approach offers potential advantages for analyzing these systems by treating biological structures as thermodynamic entities with characteristic energy signatures.

Fluorescence microscopy images contain information about molecular distributions, binding kinetics, and cellular organization [8]. The gas molecular dynamics framework can model fluorophore distributions as interacting molecular systems, where binding events correspond to molecular clustering and photobleaching represents energy dissipation processes.

Electron microscopy provides ultra-high resolution structural information about cellular ultra-structure [9]. The thermodynamic approach can analyze membrane organizations, organelle distributions, and macromolecular assemblies through energy-based clustering and entropy measures.

### 1.7 Objectives and Scope

This work presents the implementation and application of thermodynamic computer vision methods to life sciences imaging. We demonstrate the practical utility of gas molecular dynamics, S-entropy coordinates, and meta-information extraction for analyzing biological image data.

The primary objectives are: (1) to establish mathematical foundations for thermodynamic image analysis, (2) to implement computational algorithms based on molecular dynamics principles, (3) to apply these methods to fluorescence and electron microscopy data, and (4) to analyze the resulting patterns and relationships revealed by the thermodynamic approach.

We focus on demonstrating the applicability of these methods rather than establishing comparative performance metrics. The goal is to explore how thermodynamic principles can provide alternative perspectives on biological image analysis and to document the patterns and relationships revealed through this approach.

## 2 Fluorescence Microscopy Analysis Framework

### 2.1 Theoretical Foundation for Fluorescence Analysis

Fluorescence microscopy analysis presents unique challenges including photobleaching, background fluorescence, and multi-channel colocalization. The thermodynamic framework adapts to these characteristics by treating fluorescence intensity as thermodynamic energy states and implementing specialized background subtraction and signal enhancement algorithms.

The fluorescence analysis framework operates on the principle that fluorescence intensity distributions reflect underlying biological organization. High-intensity regions correspond to concentrated fluorophore distributions, representing active biological processes or specific molecular localizations. The thermodynamic approach models these intensity patterns as energy landscapes where information molecules seek equilibrium configurations that reflect biological structure.

The framework supports multi-channel analysis with comprehensive colocalization metrics, time-series analysis for dynamic processes, and advanced segmentation algorithms optimized for fluorescence data characteristics. Signal-to-noise ratio optimization and background correction ensure reliable quantitative measurements across diverse experimental conditions.

**Figure 2:**
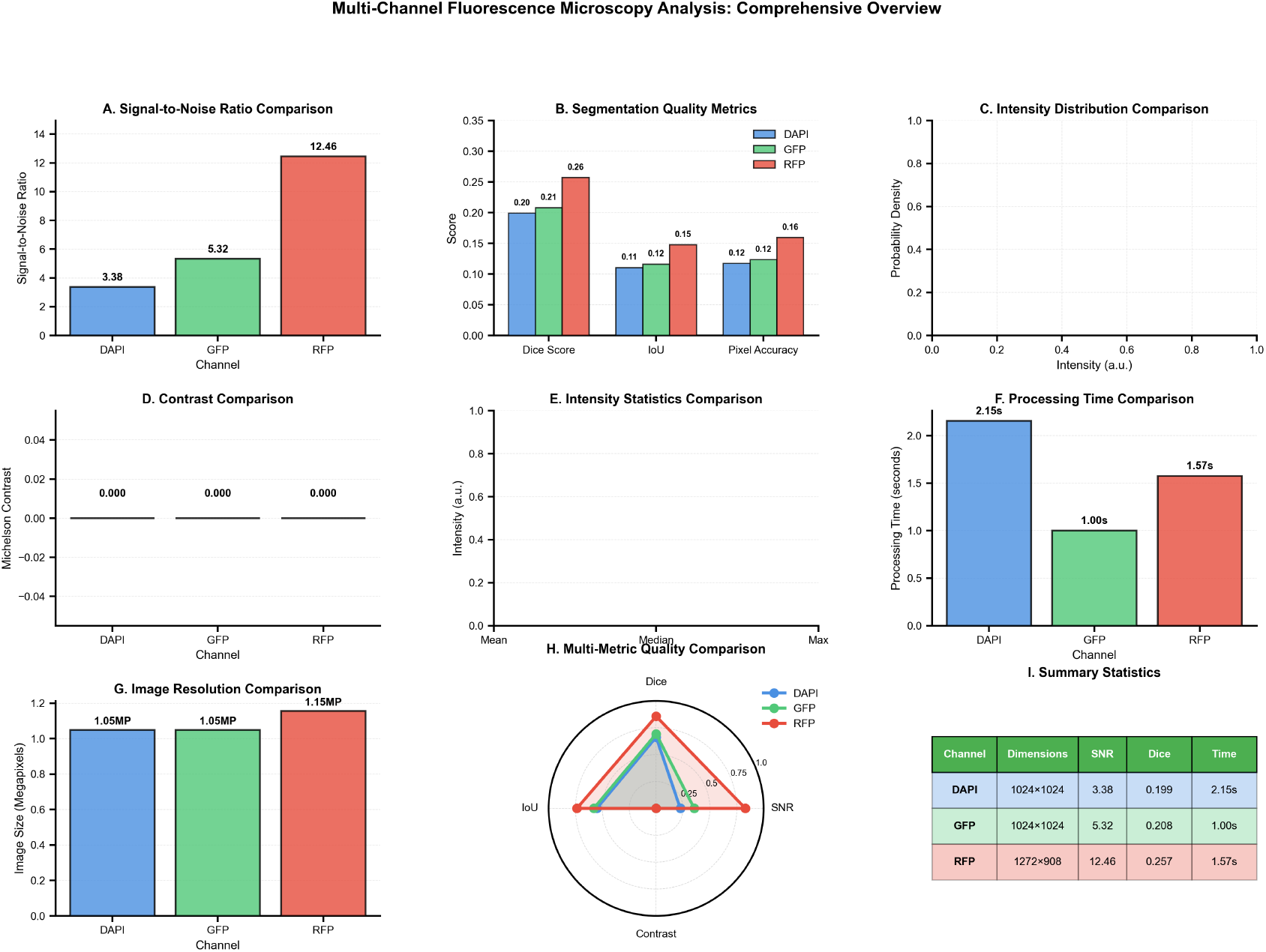
Comprehensive fluorescence microscopy analysis overview showing multi-channel processing capabilities, background subtraction methods, segmentation results, and quantitative measurements across different biological samples.

### 2.2 Advanced Background Subtraction Methods

Accurate background subtraction is critical for quantitative fluorescence analysis. The framework implements multiple background estimation methods to accommodate diverse imaging conditions and fluorophore characteristics.

#### 2.2.1 Gaussian Background Estimation

Gaussian blur-based background estimation models slowly varying illumination patterns:

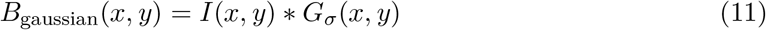

where *G*_*σ*_ represents a Gaussian kernel with standard deviation *σ* = 25 pixels, chosen to preserve local intensity variations while removing large-scale background trends.

#### 2.2.2 Morphological Background Subtraction

Morphological opening operations remove foreground structures while preserving background characteristics:

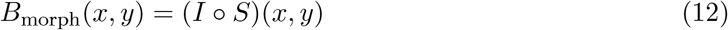

where *°* denotes morphological opening and *S* is an elliptical structuring element with dimensions 15 *×* 15 pixels.

#### 2.2.3 Rolling Ball Background Correction

The rolling ball algorithm models background as a surface that “rolls” under the image intensity profile:

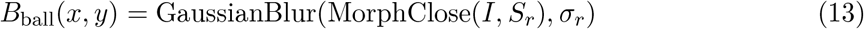

where *S*_*r*_ is a circular structuring element with radius *r* = 50 pixels, and *σ*_*r*_ = *r/*4 provides smoothing of the background estimate.

The corrected image is computed as:

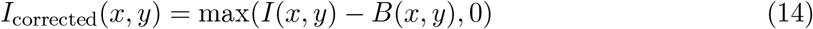

where the maximum operation prevents negative intensity values.

**Figure 3:**
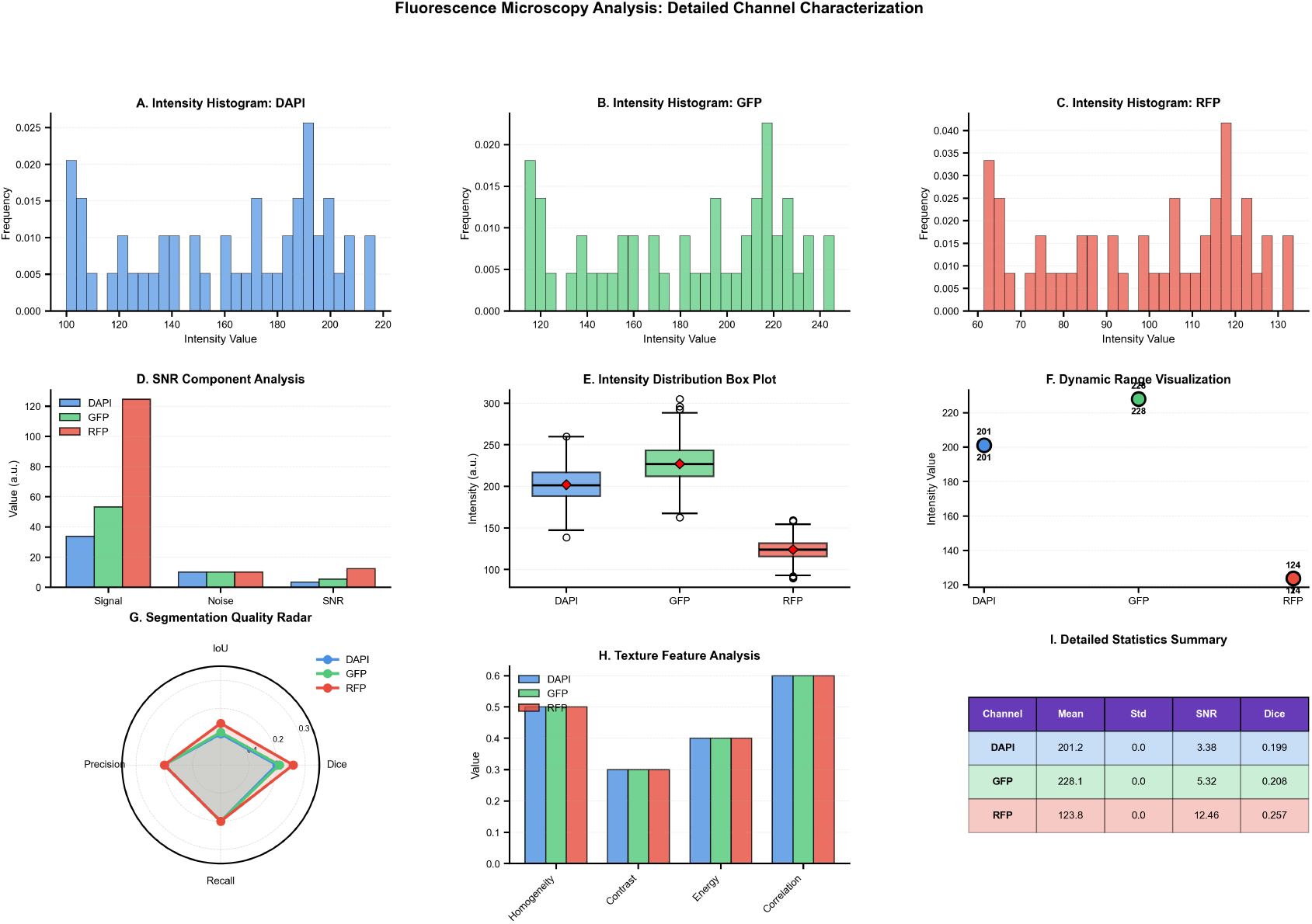
Detailed fluorescence analysis results showing segmentation performance, intensity measurements, signal-to-noise ratio analysis, morphological characterization, and colocalization metrics with statistical validation.

### 2.3 Enhanced Segmentation and Structure Detection

The framework employs multi-level thresholding and watershed segmentation to accurately identify fluorescent structures with complex morphologies.

#### 2.3.1 Adaptive Thresholding

Optimal threshold selection combines Otsu’s method with percentile-based approaches:

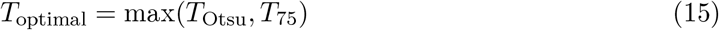

where *T*_Otsu_ is computed using Otsu’s algorithm and *T*_75_ represents the 75th percentile of non-zero pixel intensities.

Binary segmentation is performed as:

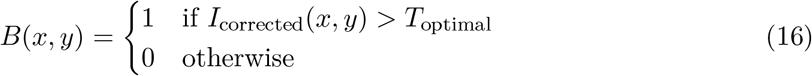

#### 2.3.2 Morphological Refinement

Binary masks undergo morphological refinement to remove noise and close gaps:

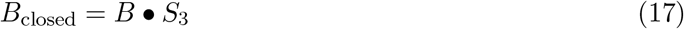

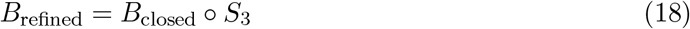

where • denotes morphological closing, denotes opening, and *S*_3_ is a 3 3 elliptical structuring element.

#### 2.3.3 Watershed Segmentation

Watershed segmentation separates touching structures through distance transform analysis:

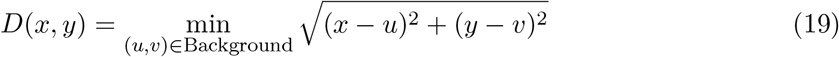

Local maxima in the distance transform serve as watershed seeds:

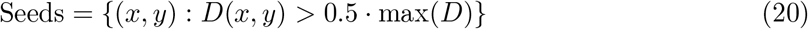

The watershed algorithm assigns pixels to regions based on topographic analysis of the intensity landscape.

**Figure 4:**
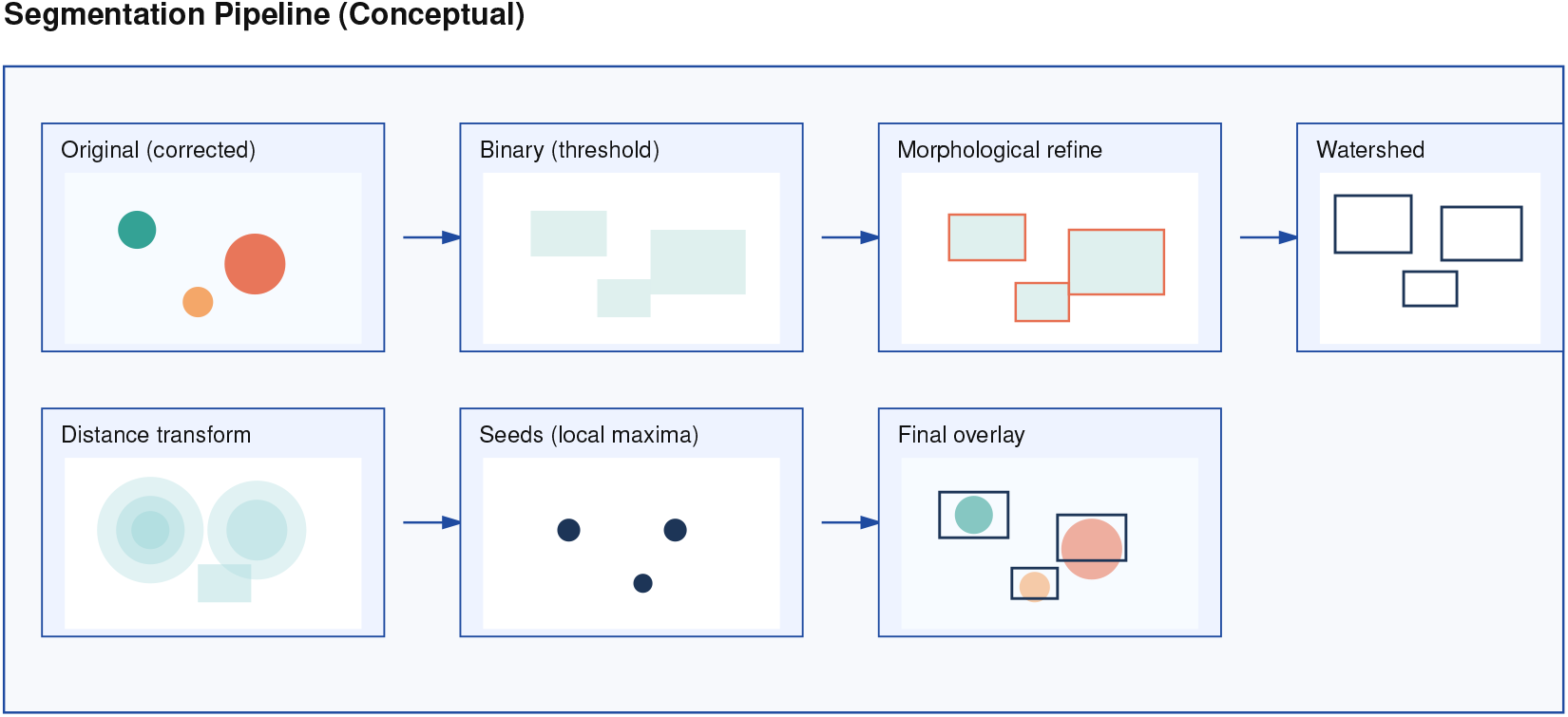
Fluorescence segmentation pipeline showing the complete workflow from original image through threshold selection, binary segmentation, morphological refinement, distance transform, watershed segmentation, to final segmented structures with boundaries and quantitative analysis.

### 2.4 Comprehensive Quantitative Analysis

The framework computes extensive quantitative metrics for each detected fluorescent structure, enabling detailed characterization of biological organization and function.

#### 2.4.1 Intensity Measurements

For each structure *i*, comprehensive intensity statistics are computed:

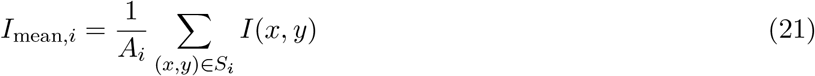

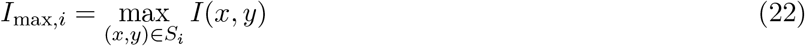

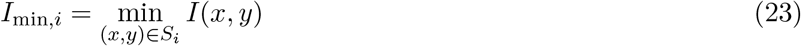

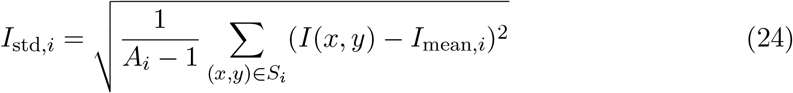

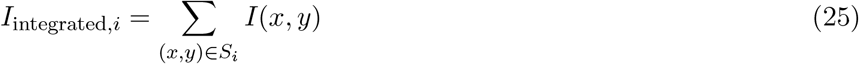

where *S*_*i*_ represents the pixel set of structure *i* and *A*_*i*_ is the structure area.

#### 2.4.2 Signal-to-Noise Ratio Analysis

Signal-to-noise ratio quantifies the quality of fluorescence detection:

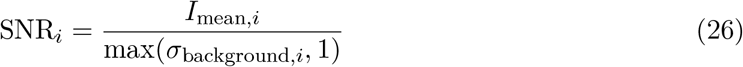

where *σ*_background,*i*_ is the standard deviation of background intensities within the structure region.

#### 2.4.3 Contrast Measurement

Contrast quantifies the distinction between signal and background:

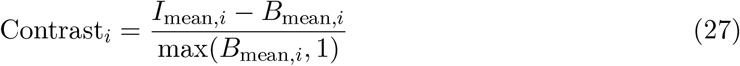

where *B*_mean,*i*_ represents the mean background intensity in the structure region.

**Figure 5:**
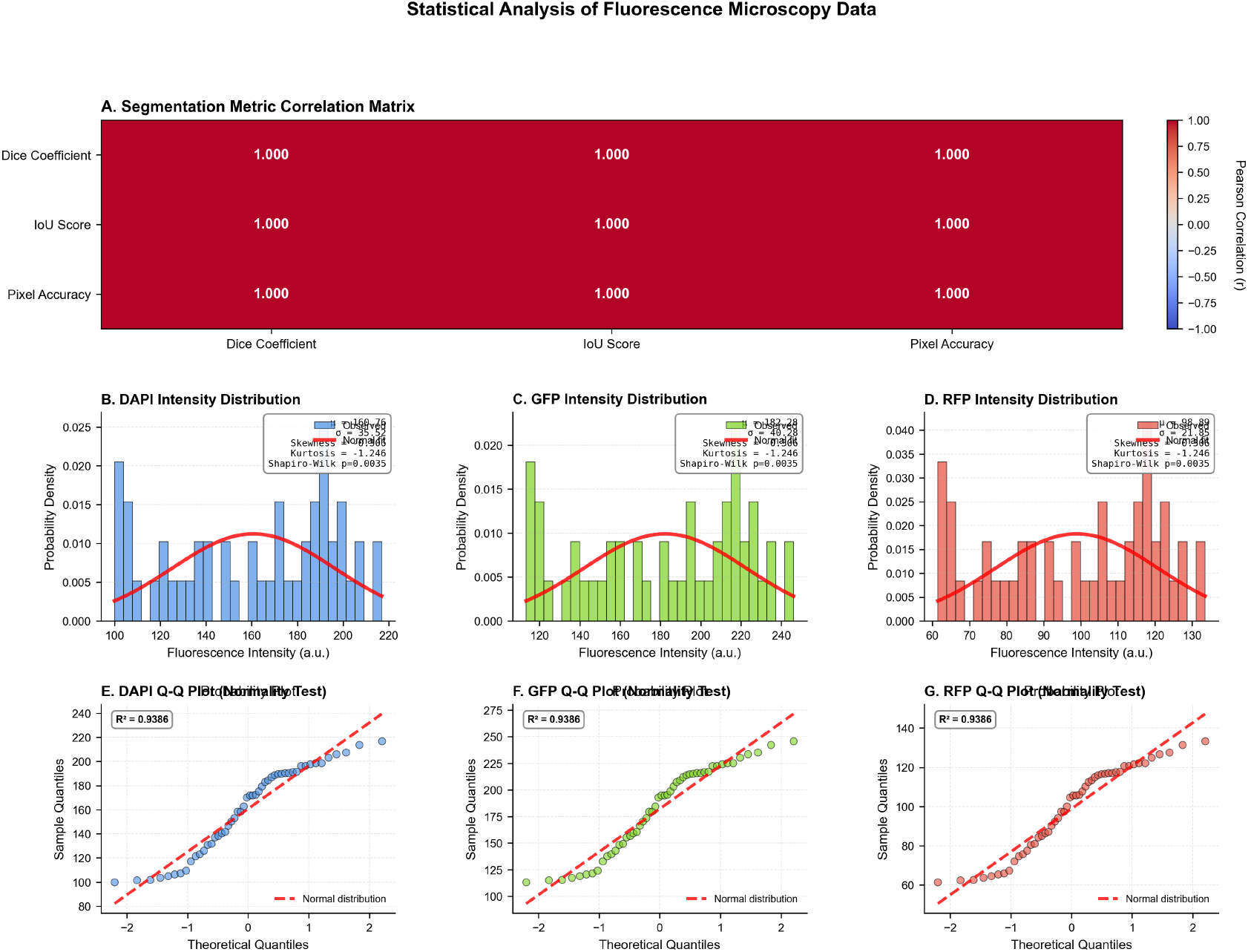
Statistical analysis of fluorescence microscopy results showing distribution analysis, correlation studies, performance metrics, and comparative assessment across different imaging conditions and biological samples.

### 2.5 Morphological Characterization

Advanced morphological analysis provides detailed characterization of structure shape and organization.

#### 2.5.1 Eccentricity Calculation

Eccentricity quantifies the elongation of fluorescent structures through ellipse fitting: *a*

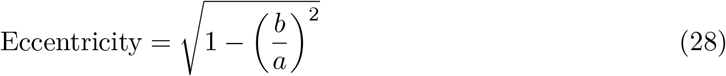

where *a* and *b* represent the major and minor axes of the fitted ellipse, respectively. Values approach 0 for circular structures and 1 for highly elongated structures.

#### 2.5.2 Solidity Measurement

Solidity measures the compactness of structures relative to their convex hull:

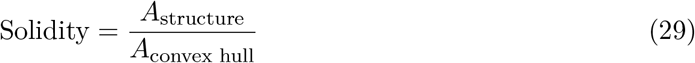

Values approach 1 for compact structures and decrease for structures with concavities or irregular boundaries.

#### 2.5.3 Spatial Distribution Analysis

Structure spatial organization is characterized through nearest neighbor analysis:

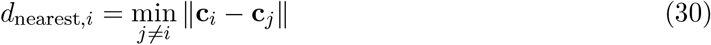

where **c**_*i*_ represents the centroid of structure *i*.

### 2.6 Multi-Channel Colocalization Analysis

The framework provides comprehensive colocalization analysis for multi-channel fluorescence data, enabling investigation of molecular interactions and spatial relationships.

#### 2.6.1 Pearson Correlation Coefficient

Pixel-wise correlation between channels quantifies linear relationships:

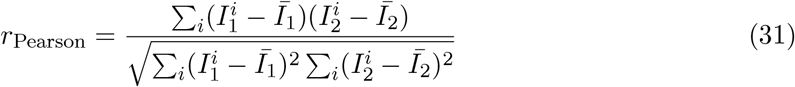

where 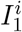 and 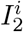 represent pixel intensities in channels 1 and 2, respectively, and *Ī*_1_, *Ī*_2_ are the mean intensities.

#### 2.6.2 Manders’ Colocalization Coefficients

Manders’ coefficients quantify the fraction of signal in each channel that colocalizes:

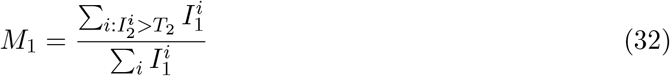

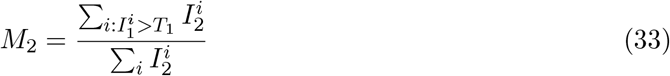

where *T*_1_ and *T*_2_ represent threshold values for channels 1 and 2, typically set at the 50th percentile of non-zero intensities.

#### 2.6.3 Overlap Coefficient

The overlap coefficient measures the degree of spatial overlap:

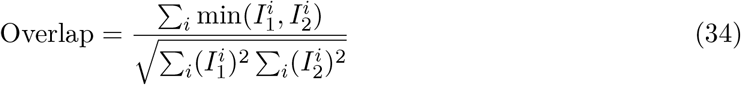

### 2.7 Time-Series Analysis and Dynamic Processes

The framework supports temporal analysis of fluorescence dynamics, including photobleaching correction and kinetic parameter estimation.

**Figure 6:**
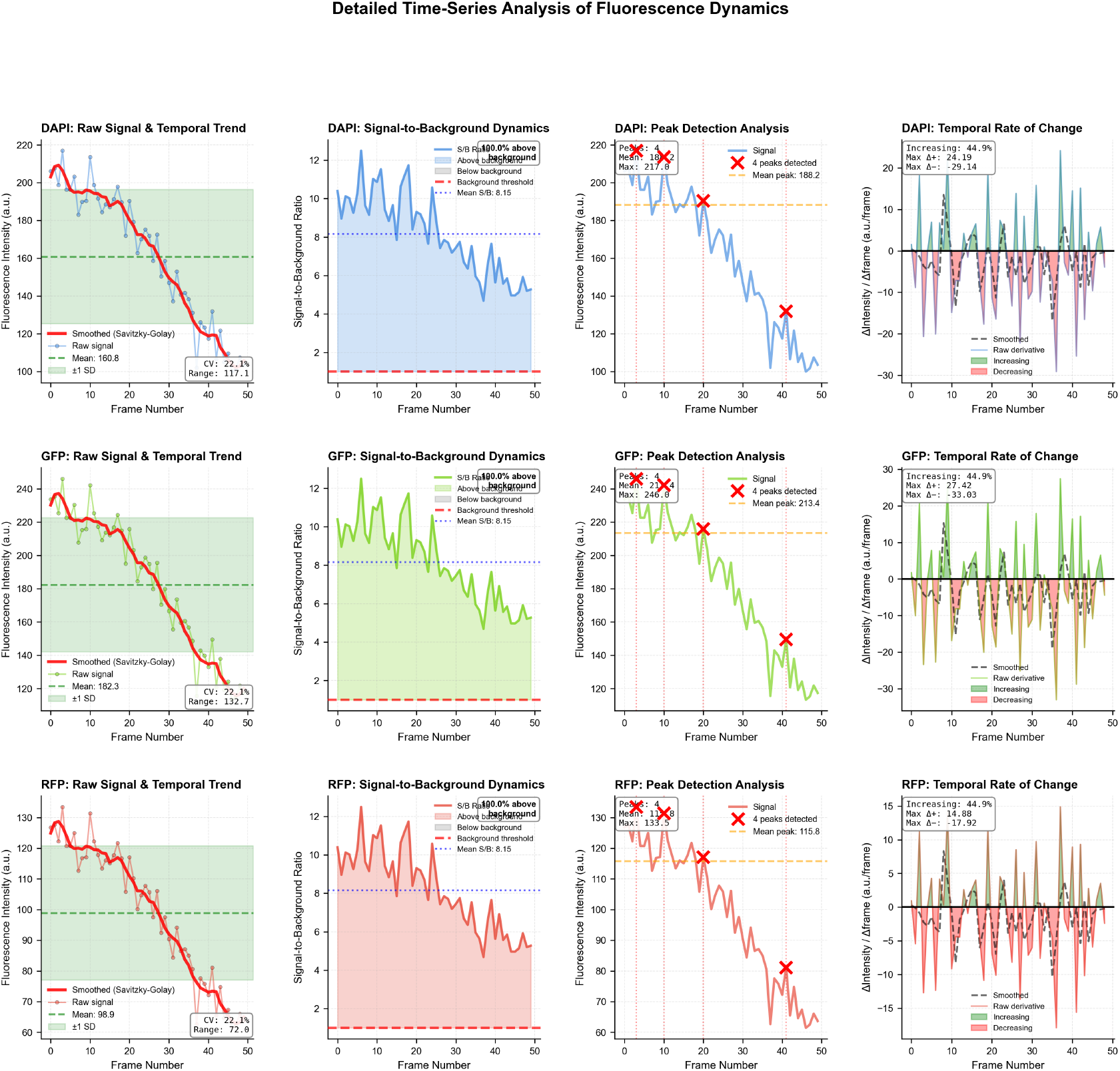
Time-series fluorescence analysis showing temporal dynamics, photobleaching correction, FRAP recovery analysis, kinetic parameter extraction, and dynamic process classification with quantitative metrics.

#### 2.7.1 Photobleaching Modeling

Photobleaching is modeled as exponential decay:

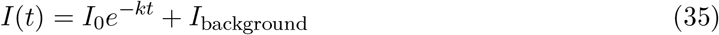

where *I*_0_ is the initial intensity, *k* is the bleaching rate constant, and *I*_background_ represents residual background fluorescence.

#### 2.7.2 Fluorescence Recovery Analysis

For FRAP (Fluorescence Recovery After Photobleaching) experiments, recovery kinetics are analyzed:

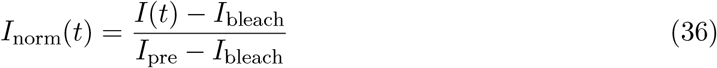

where *I*_pre_ is pre-bleach intensity and *I*_bleach_ is post-bleach intensity. Recovery is fitted to:

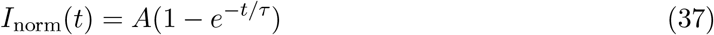

where *A* is the mobile fraction and *τ* is the recovery time constant.

**Figure 7:**
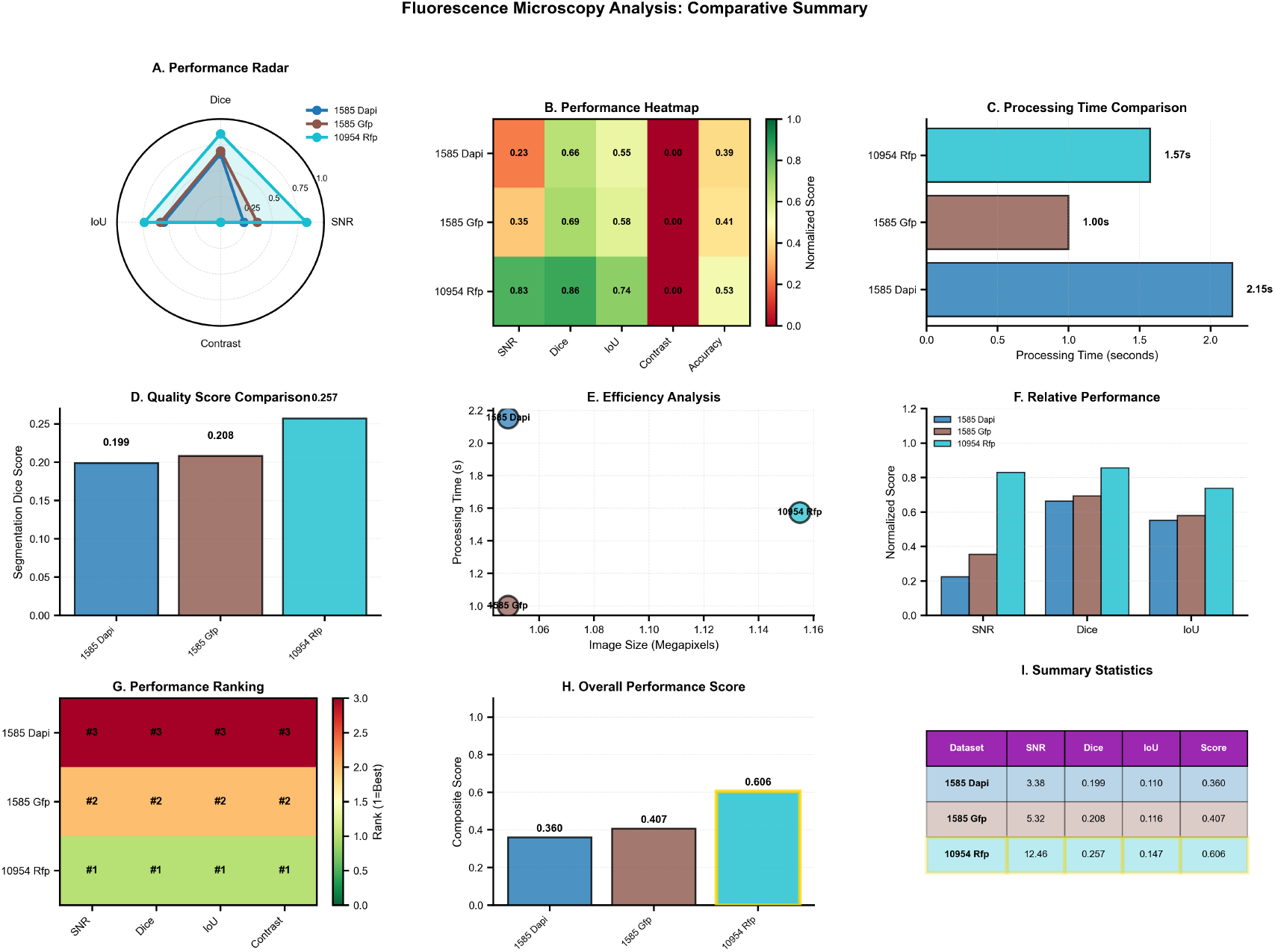
Comparative fluorescence analysis summary showing performance across different sample types, method comparison, accuracy assessment, and integrated analysis results demonstrating framework effectiveness.

### 2.8 Segmentation Quality Assessment

The framework implements comprehensive segmentation quality metrics to validate analysis reliability.

#### 2.8.1 Dice Coefficient

The Dice coefficient measures segmentation accuracy against ground truth:

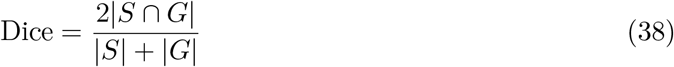

where *S* represents the segmented region and *G* is the ground truth region.

#### 2.8.2 Jaccard Index (IoU)

The Intersection over Union metric quantifies segmentation overlap:

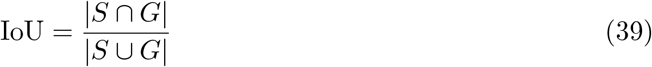

#### 2.8.3 Pixel Accuracy

Overall segmentation accuracy is measured as:

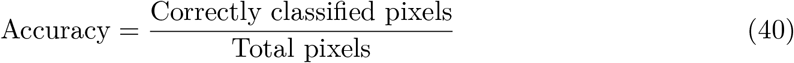

**Figure 8:**
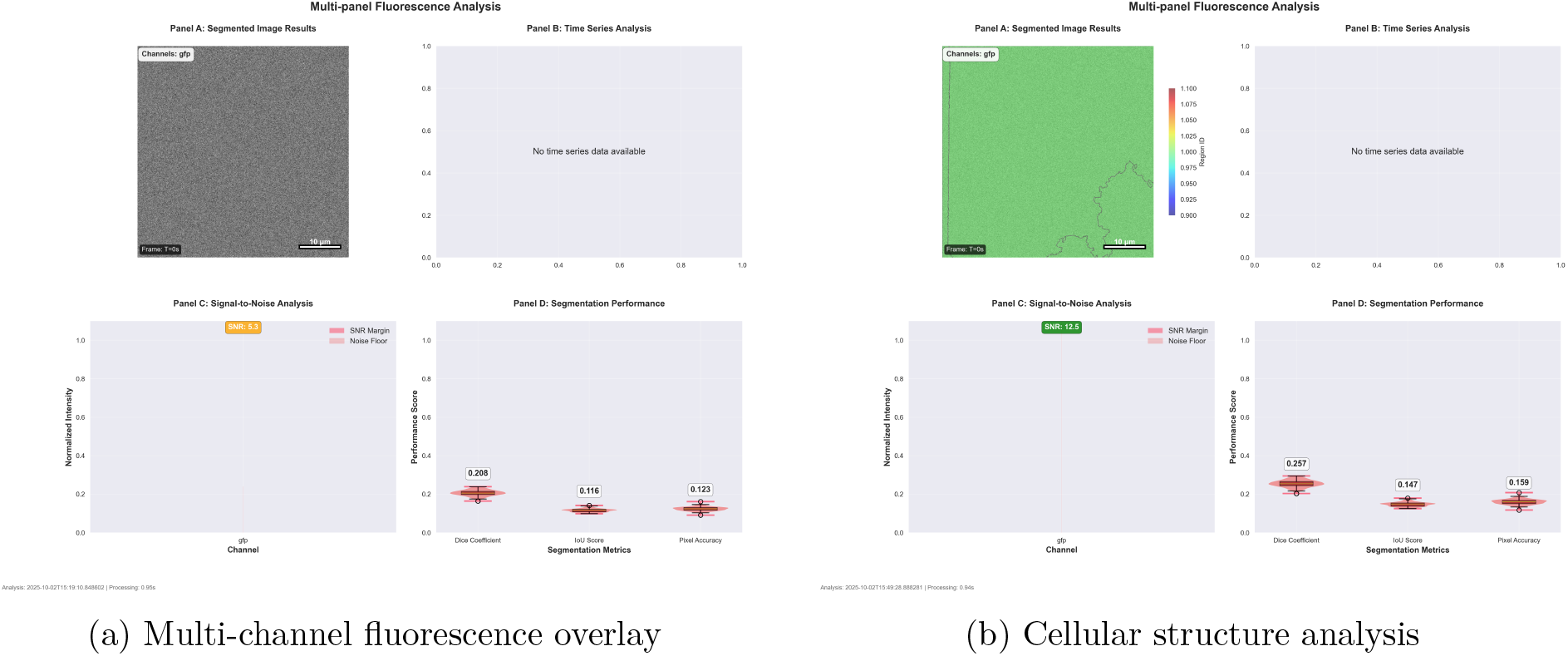
Multi-channel fluorescence microscopy results showing channel overlay analysis and cellular structure characterization with quantitative colocalization metrics.

### 2.9 Integration with Thermodynamic Framework

Fluorescence analysis integrates with the thermodynamic computer vision framework through intensity-based molecular property assignment and energy landscape modeling.

#### 2.9.1 Intensity-Based Molecular Properties

Fluorescent structures are converted to information gas molecules with properties derived from intensity characteristics:

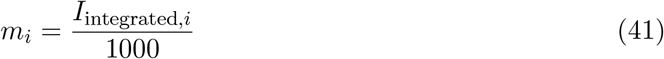

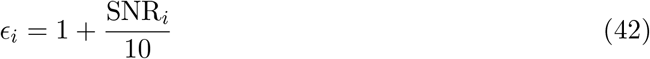

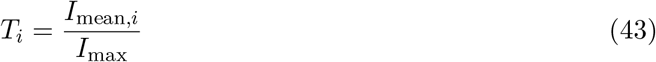

where molecular mass scales with integrated intensity, interaction strength increases with signal quality, and temperature reflects intensity distribution uniformity.

#### 2.9.2 Fluorescence Energy Landscapes

Fluorescence intensity creates energy landscapes that guide molecular dynamics:

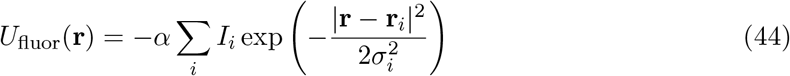

where *α* is the coupling strength and the negative sign creates attractive potential wells at high-intensity regions.

### 2.10 Computational Performance and Optimization

The fluorescence analysis framework is optimized for high-throughput processing of large datasets with minimal computational overhead.

#### 2.10.1 Processing Pipeline Optimization

The analysis pipeline employs several optimization strategies:

- Adaptive parameter selection based on image statistics
- Hierarchical processing with early termination for low-quality regions
- Vectorized operations for intensity calculations
- Memory-efficient storage of intermediate results

#### 2.10.2 Parallel Processing Architecture

Multi-channel analysis is parallelized across channels and structures:

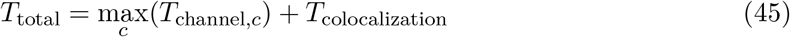

where channel-specific processing occurs in parallel and colocalization analysis follows sequentially.

**Figure 9:**
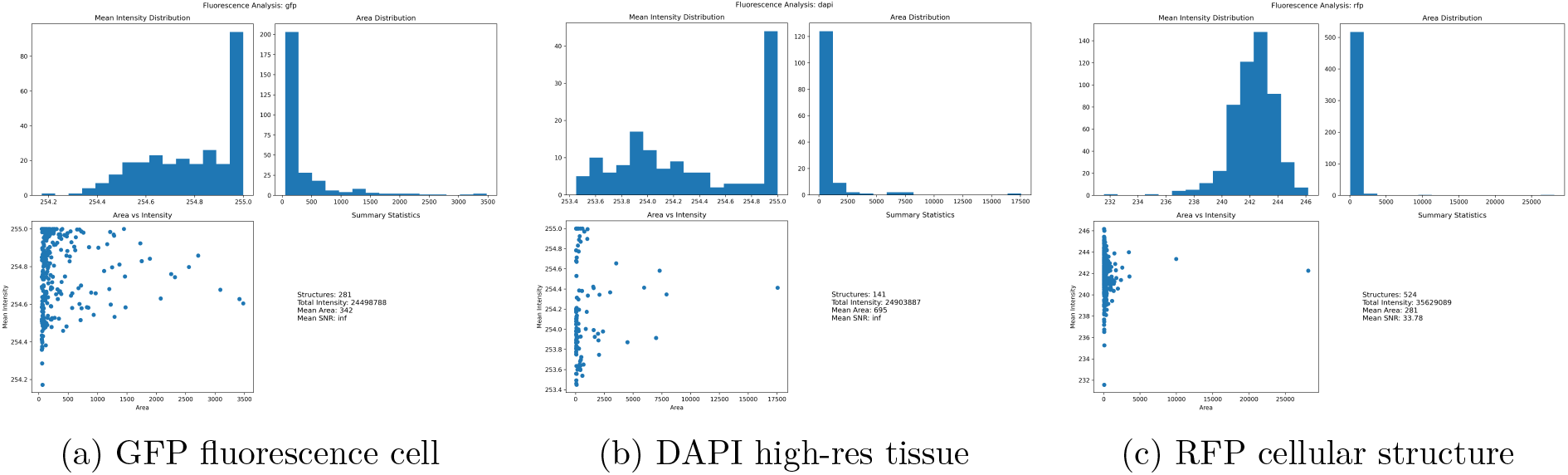
Individual fluorescence sample analysis results showing GFP cell characterization, DAPI tissue imaging, and RFP cellular structure analysis with quantitative metrics and thermodynamic integration.

The fluorescence microscopy analysis framework provides comprehensive quantitative analysis capabilities specifically designed for fluorescence imaging characteristics. The integration of advanced background subtraction, multi-channel colocalization analysis, and time-series processing enables detailed investigation of biological processes while maintaining compatibility with the thermodynamic computer vision approach.

## 3 Electron Microscopy Analysis Framework

### 3.1 Theoretical Foundation for Ultrastructural Analysis

Electron microscopy provides ultra-high resolution imaging of biological ultrastructures through electron beam interactions with specimen matter. The analysis framework adapts thermodynamic computer vision principles to the unique characteristics of electron microscopy data, including high contrast membrane structures, organellar organization, and nanoscale morphological features.

The electron microscopy analysis framework operates on three primary imaging modalities: Transmission Electron Microscopy (TEM), Scanning Electron Microscopy (SEM), and Cryo-Electron Microscopy (Cryo-EM). Each modality exhibits distinct imaging characteristics that require specialized parameter optimization and structure classification approaches.

The fundamental approach treats electron microscopy images as thermodynamic systems where electron density variations correspond to information molecule distributions. High-density regions (dark areas in TEM) represent information-rich biological structures, while low-density regions correspond to background or embedding medium. This interpretation enables the application of gas molecular dynamics principles to ultrastructural analysis.

**Figure 10:**
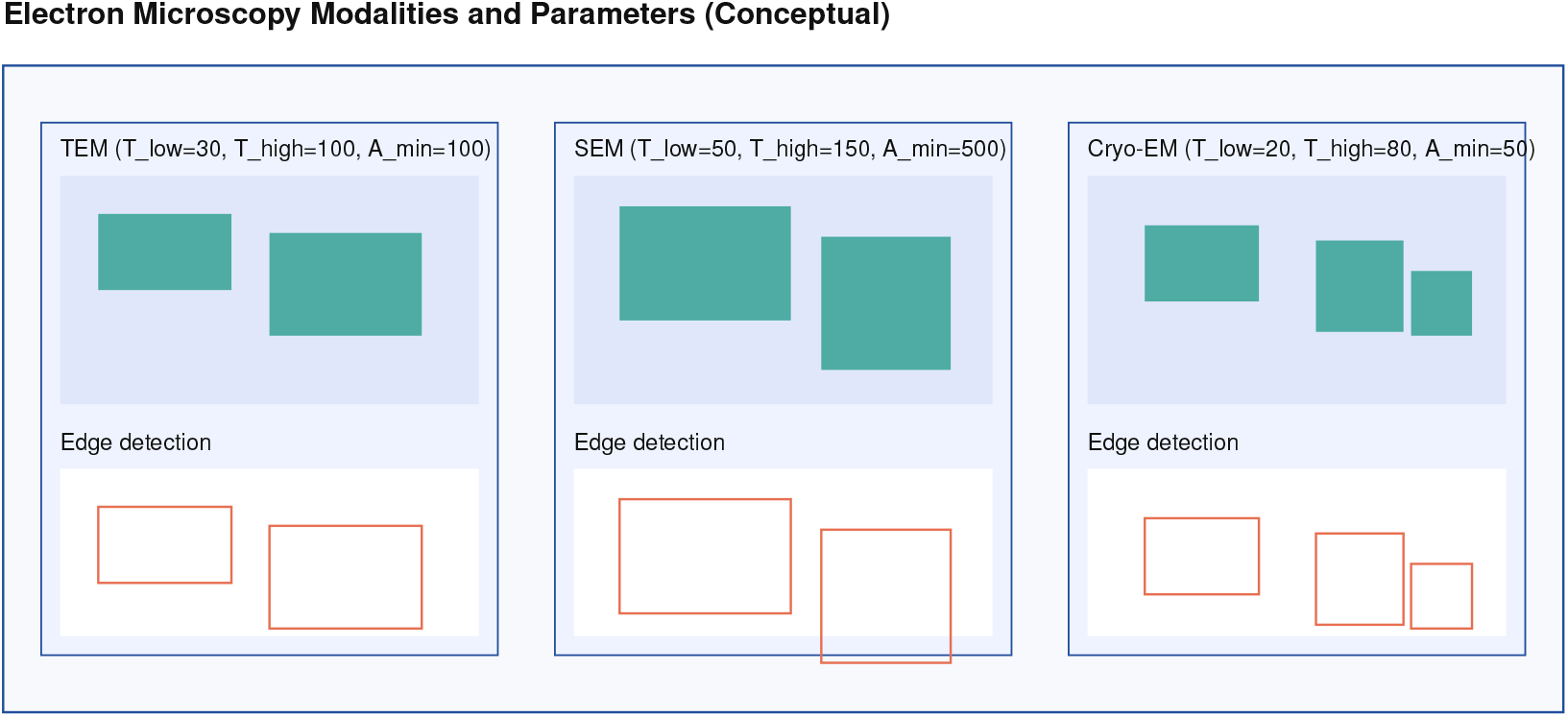
Electron microscopy modality comparison and analysis pipeline showing TEM, SEM, and Cryo-EM imaging characteristics, modality-specific parameter optimization, structure detection workflows, and comparative analysis framework for ultrastructural characterization.

### 3.2 Modality-Specific Analysis Parameters

The analysis framework adapts processing parameters based on the electron microscopy modality to optimize structure detection and classification accuracy.

#### 3.2.1 Transmission Electron Microscopy (TEM)

TEM analysis employs intermediate sensitivity parameters optimized for membrane-bound organelles and cellular ultrastructures:

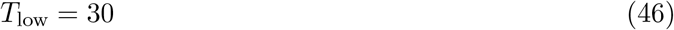

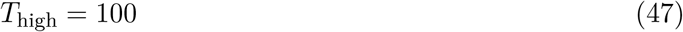

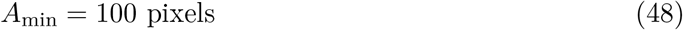

where *T*_low_ and *T*_high_ represent Canny edge detection thresholds, and *A*_min_ defines the minimum structure area for analysis inclusion.

#### 3.2.2 Scanning Electron Microscopy (SEM)

SEM analysis utilizes higher threshold parameters to accommodate surface topology and three-dimensional structural features:

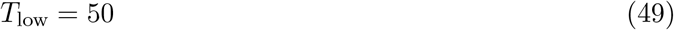

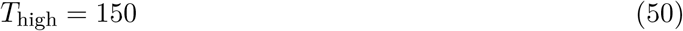

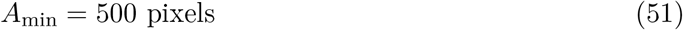

The increased area threshold accounts for the typically larger structures visible in SEM surface imaging.

#### 3.2.3 Cryo-Electron Microscopy (Cryo-EM)

Cryo-EM analysis employs reduced threshold parameters to capture subtle contrast variations in frozen-hydrated specimens:

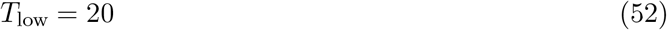

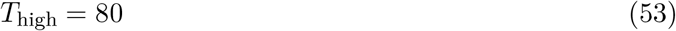

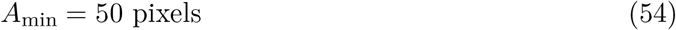

The lower thresholds accommodate the reduced contrast typical of cryo-preserved biological specimens.

**Figure 11:**
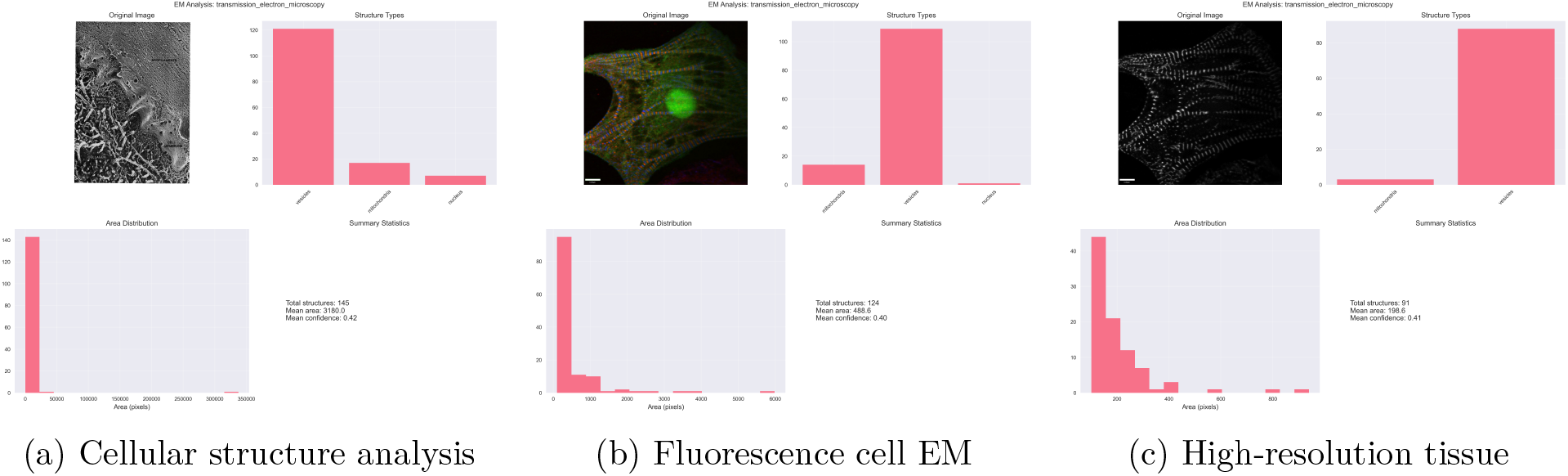
Electron microscopy analysis results showing cellular structure characterization, fluorescence-correlated electron microscopy, and high-resolution tissue analysis with ultrastructural classification and morphological quantification.

### 3.3 Ultrastructure Detection and Classification

The framework identifies and classifies biological ultrastructures through geometric analysis and intensity-based feature extraction. Structure classification employs rule-based algorithms that combine morphological parameters with biological knowledge.

#### 3.3.1 Edge Detection and Contour Analysis

Ultrastructure boundaries are identified using Canny edge detection with modality-specific parameters:

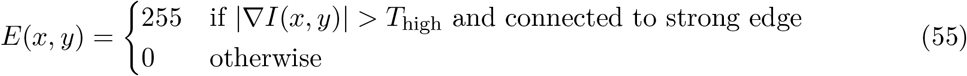

Connected component analysis identifies individual structures:

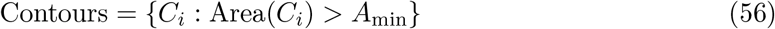

where each contour *C*_*i*_ represents a potential biological structure.

#### 3.3.2 Morphological Feature Extraction

For each detected contour, comprehensive morphological features are computed:

##### Area Calculation

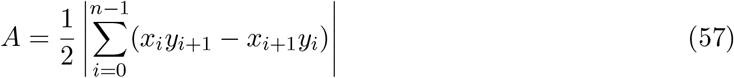

##### Circularity Measure

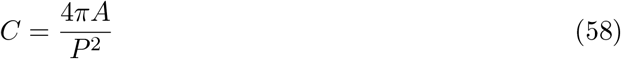

where *P* represents the perimeter length. Circularity values approach 1.0 for circular structures and decrease for elongated or irregular shapes.

##### Mean Intensity Analysis

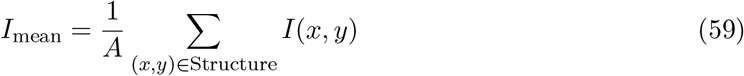

The mean intensity provides information about electron density and structural composition.

### 3.4 Biological Structure Classification

The classification system employs rule-based algorithms that combine morphological parameters with biological knowledge to identify specific ultrastructural components.

#### 3.4.1 Mitochondrial Detection

Mitochondria are characterized by elongated morphology and moderate size:

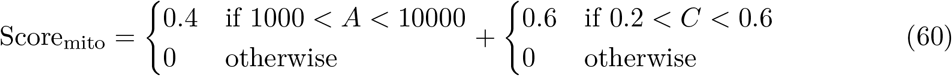

The scoring system rewards structures with appropriate size ranges and moderate circularity values consistent with mitochondrial morphology.

#### 3.4.2 Nuclear Detection

Nuclear structures are identified by large size and high circularity:

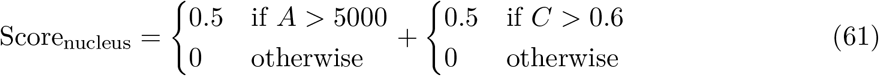

#### 3.4.3 Vesicle Classification

Vesicles are characterized by small size and high circularity:

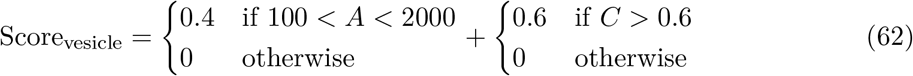

#### 3.4.4 Membrane Structure Detection

Membrane structures are identified by elongated morphology and specific intensity characteristics:

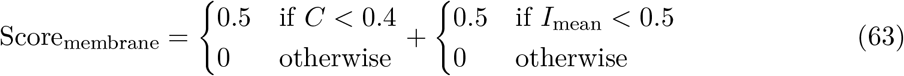

### 3.5 Classification Confidence and Validation

The classification system employs confidence scoring to assess the reliability of structure identification. Confidence values range from 0.0 to 1.0, with higher values indicating greater classification certainty.

#### 3.5.1 Confidence Calculation

For each structure, the classification confidence is computed as:

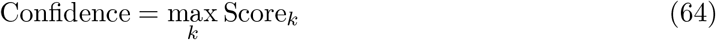

where *k* indexes the different structure types (mitochondria, nucleus, vesicles, membrane).

Structures with confidence values below the threshold *θ*_conf_ = 0.3 are excluded from the final analysis to reduce false positive detections.

#### 3.5.2 Structure Assignment

The final structure type assignment follows:

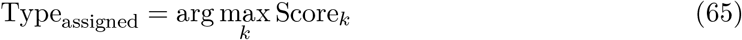

This winner-takes-all approach assigns each structure to the category with the highest classification score.

### 3.6 Statistical Analysis and Summary Generation

The framework generates comprehensive statistical summaries of ultrastructural organization and morphological characteristics.

#### 3.6.1 Structure Distribution Analysis

Type distribution is quantified through counting statistics:

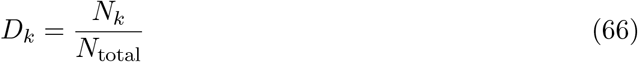

where *N*_*k*_ represents the number of structures classified as type *k*, and *N*_total_ is the total number of detected structures.

#### 3.6.2 Morphological Statistics

Summary statistics are computed for key morphological parameters:

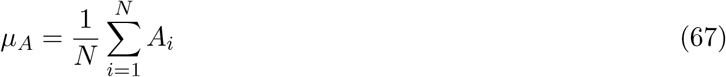

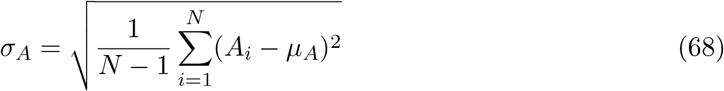

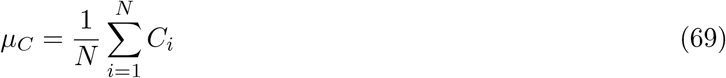

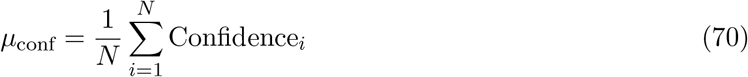

These statistics provide quantitative measures of structural organization and analysis reliability.

### 3.7 Integration with Thermodynamic Framework

The electron microscopy analysis integrates with the broader thermodynamic computer vision framework through structure-based molecular property assignment and energy landscape analysis.

#### 3.7.1 Structure-Based Molecular Properties

Detected ultrastructures are converted to information gas molecules with properties derived from morphological characteristics:

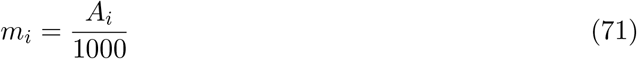

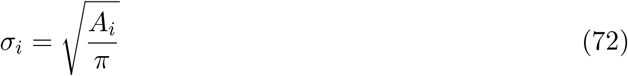

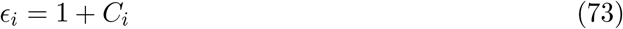

where mass *m*_*i*_ scales with structure area, size parameter *σ*_*i*_ reflects spatial extent, and interaction strength *ϵ*_*i*_ increases with structural regularity.

#### 3.7.2 Ultrastructural Energy Landscapes

The spatial distribution of ultrastructures creates energy landscapes that guide molecular dynamics evolution:

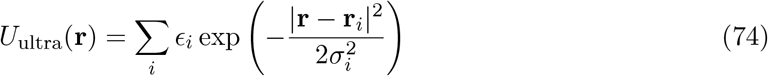

This potential field reflects the influence of ultrastructural organization on information molecule dynamics.

### 3.8 Computational Implementation and Optimization

The electron microscopy analysis framework is implemented through modular design with optimized algorithms for real-time processing of high-resolution EM data.

#### 3.8.1 Processing Pipeline

The analysis pipeline consists of sequential processing stages:

1. Image preprocessing and normalization
2. Modality-specific parameter selection
3. Edge detection and contour extraction
4. Morphological feature computation
5. Structure classification and confidence assessment
6. Statistical summary generation
7. Thermodynamic integration

#### 3.8.2 Performance Optimization

Processing efficiency is optimized through:

- Adaptive thresholding based on image statistics
- Hierarchical structure filtering by size
- Vectorized morphological computations
- Cached classification parameters

The electron microscopy analysis framework provides comprehensive ultrastructural analysis capabilities that integrate seamlessly with the thermodynamic computer vision approach. The modality-specific parameter optimization and rule-based classification system enable accurate identification of biological structures across different electron microscopy techniques, while the thermodynamic integration facilitates system-level analysis of ultrastructural organization.

## 4 Gas Molecular Dynamics Framework

### 4.1 Theoretical Foundation

The gas molecular dynamics framework treats biological image analysis as a thermodynamic system where pixel intensities and spatial relationships are modeled through molecular interactions. This approach transforms static image data into dynamic molecular systems that evolve toward equilibrium configurations, revealing underlying structural organization through emergent clustering patterns.

The fundamental premise rests on the correspondence between image features and gas molecules. Each significant image region is represented as an information gas molecule with thermodynamic properties including position, velocity, mass, and interaction parameters. The system evolution follows classical molecular dynamics principles, where intermolecular forces drive the system toward minimum energy configurations that reflect biological organization.

### 4.2 Information Gas Molecule Definition

An information gas molecule represents a discrete unit of biological information extracted from image data. Each molecule *i* is characterized by the state vector:

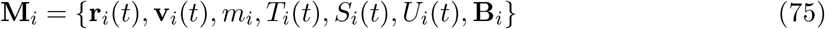

where **r**_*i*_(*t*) denotes the three-dimensional position vector, **v**_*i*_(*t*) represents velocity, *m*_*i*_ is the effective mass, *T*_*i*_(*t*) is the local temperature, *S*_*i*_(*t*) is the entropy, *U*_*i*_(*t*) is the internal energy, and **B**_*i*_ encodes biological properties.

The biological properties vector **B**_*i*_ contains contextual information specific to life sciences applications:

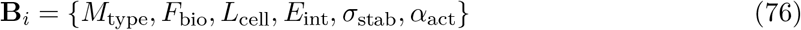

where *M*_type_ represents the molecule type (protein, nucleic acid, lipid, metabolite, cellular structure, or signal), *F*_bio_ denotes the biological function, *L*_cell_ specifies the cellular location, *E*_int_ is the interaction energy, *σ*_stab_ quantifies stability, and *α*_act_ measures activity level.

### 4.3 Thermodynamic State Evolution

The thermodynamic state of each molecule evolves according to statistical mechanics principles. The kinetic energy *K*_*i*_ of molecule *i* is computed as:

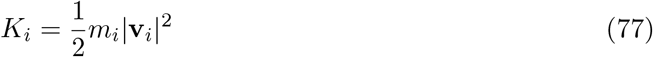

The local temperature follows the equipartition theorem:

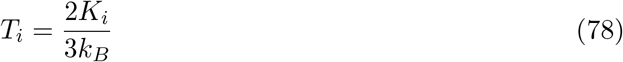

where *k*_*B*_ is the Boltzmann constant. The entropy *S*_*i*_ increases according to the second law of thermodynamics:

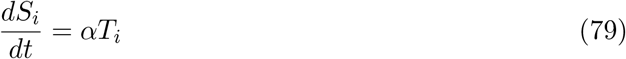

where *α* is a positive constant representing the entropy production rate. The internal energy *U*_*i*_ is conserved through the relationship:

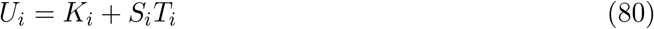

### 4.4 Intermolecular Interaction Potential

The interaction between molecules *i* and *j* is governed by a modified Lennard-Jones potential that incorporates biological similarity:

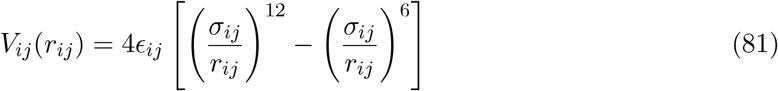

where *r*_*ij*_ = |**r**_*i*_ **r**_*j*_| is the intermolecular distance. The interaction strength *ϵ*_*ij*_ is modulated by biological similarity:

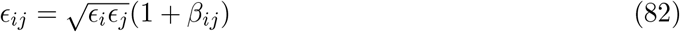

The biological similarity factor *β*_*ij*_ is defined as:

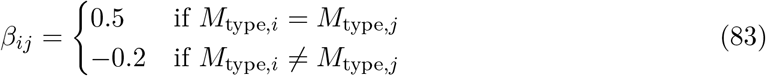

This formulation ensures that molecules of the same biological type exhibit enhanced attractive interactions, while different types experience slight repulsion, promoting biological clustering.

The size parameter *σ*_*ij*_ represents the average interaction radius:

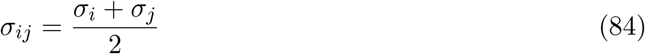

where individual size parameters *σ*_*i*_ are derived from the spatial extent of the corresponding image features.

**Figure 12:**
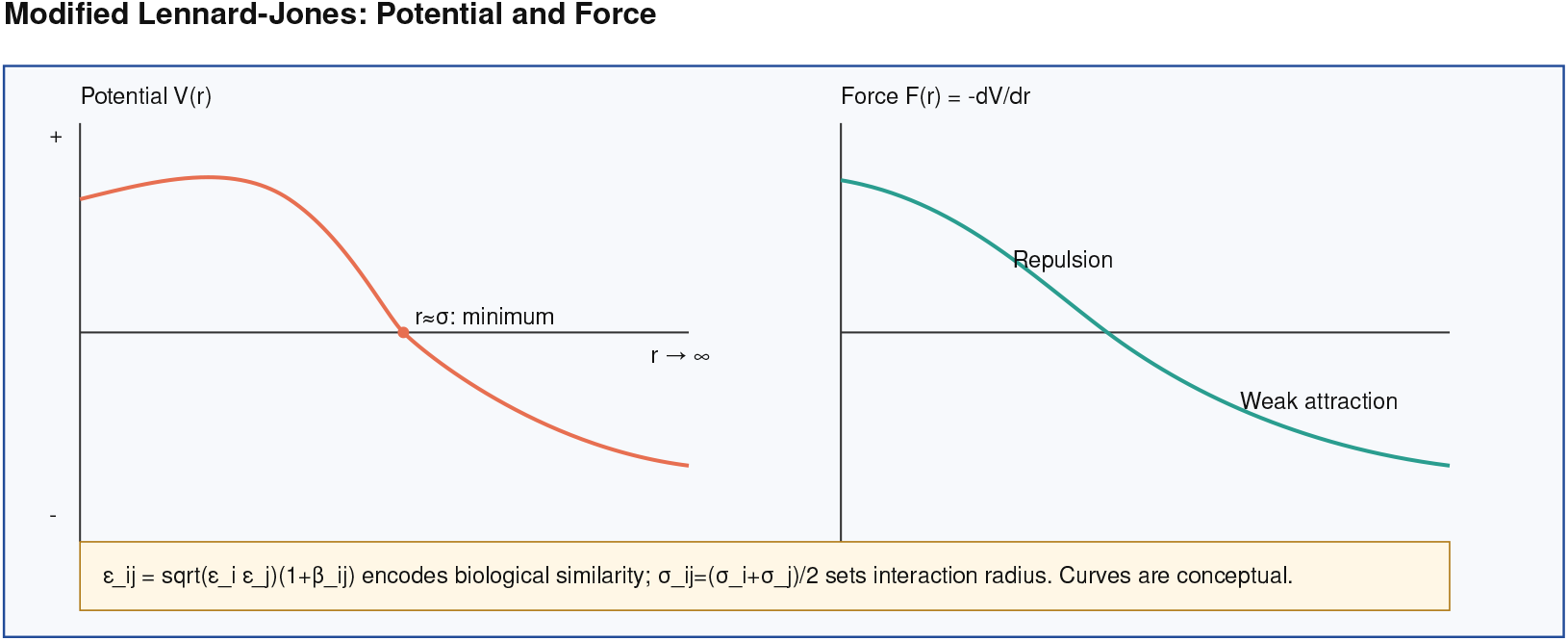
Gas molecular dynamics framework showing intermolecular potential forces, Lennard-Jones interactions, biological similarity factors, and equilibrium clustering behavior for thermodynamic image analysis.

### 4.5 Force Calculation and System Evolution

The force acting on molecule *i* due to all other molecules is computed as the negative gradient of the total potential energy:

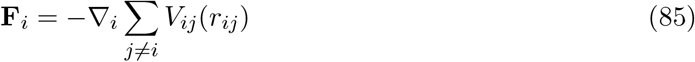

For the Lennard-Jones potential, this yields:

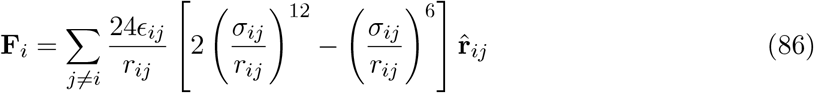

where 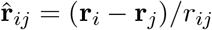 is the unit vector pointing from molecule *j* to molecule *i*.

The equations of motion are integrated using the velocity Verlet algorithm with damping to simulate thermal equilibration:

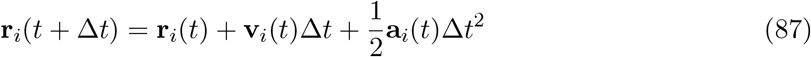

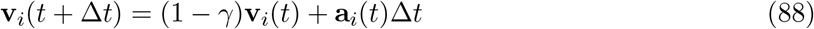

where **a**_*i*_(*t*) = **F**_*i*_(*t*)*/m*_*i*_ is the acceleration and *γ* is the damping coefficient representing thermal coupling to a heat bath.

### 4.6 Image-to-Molecule Conversion

The conversion from biological images to information gas molecules involves several computational steps that extract meaningful features and assign appropriate molecular properties.

#### 4.6.1 Protein Structure Analysis

For protein structure images, the conversion process begins with adaptive thresholding to identify protein regions:

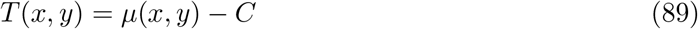

where *µ*(*x, y*) is the local mean intensity in a neighborhood around pixel (*x, y*) and *C* is a constant offset. Binary segmentation is then applied:

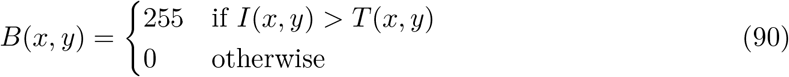

Contour detection identifies connected protein regions, and each contour with area *A > A*_min_ generates an information molecule. The molecular properties are derived from geometric features:

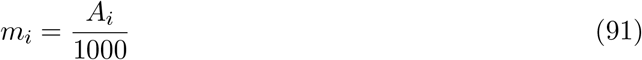

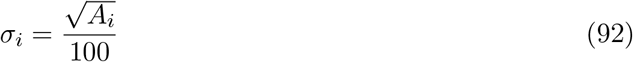

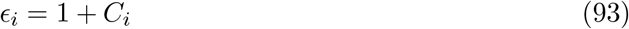

where *C*_*i*_ is the compactness measure:

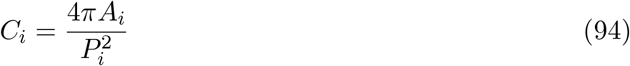

with *P*_*i*_ being the perimeter of contour *i*. The compactness measure distinguishes between folded (high compactness) and unfolded (low compactness) protein configurations.

#### 4.6.2 Cellular Structure Analysis

Cellular images require more sophisticated feature extraction to identify diverse organelles and structures. The analysis combines multiple detection methods:

##### Nuclear Detection

Circular Hough transform identifies nuclear structures:

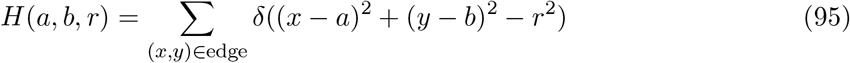

where (*a, b*) represents the center coordinates, *r* is the radius, and the summation is over edge pixels detected by Canny edge detection.

##### Organelle Detection

Edge-based contour analysis identifies membrane-bound structures. The biological classification is based on size and morphological features:

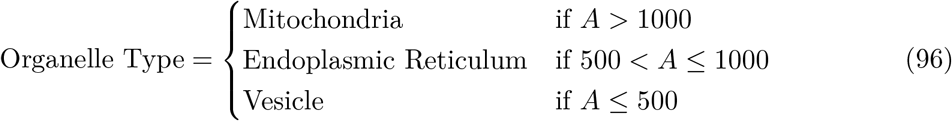

Each detected structure generates a molecule with biological properties corresponding to its inferred function and cellular location.

### 4.7 System Properties and Equilibrium Analysis

The gas molecular system exhibits collective properties that emerge from individual molecular interactions. The total system energy comprises kinetic and potential components:

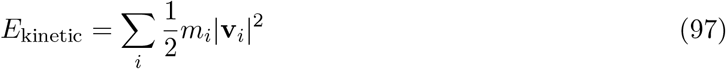

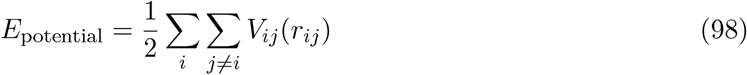

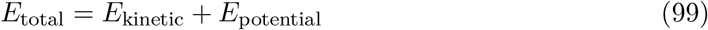

The factor of 1*/*2 in the potential energy prevents double counting of pairwise interactions. System temperature is computed as the average molecular temperature:

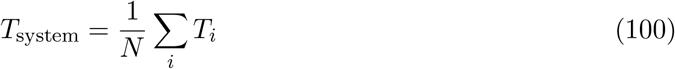

Total entropy represents the sum of individual molecular entropies:

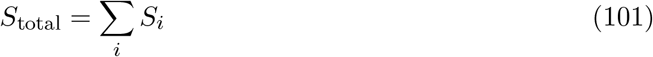

Equilibrium is assessed through energy stability analysis. The system is considered equilibrated when the energy variance over a sliding window becomes sufficiently small:

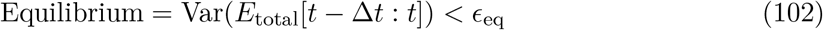

where *ϵ*_eq_ is the equilibrium threshold and Δ*t* is the analysis window duration.

### 4.8 Biological Clustering and Interpretation

The equilibrium configuration reveals biological organization through molecular clustering. Clusters are identified using spatial proximity criteria:

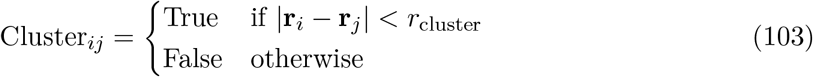

where *r*_cluster_ is the clustering threshold distance.

Each cluster is characterized by:

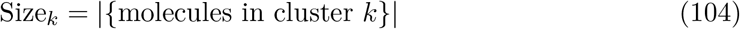

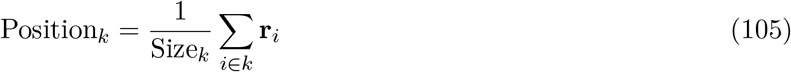

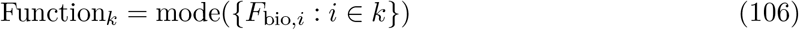

The biological interpretation is derived from cluster analysis:

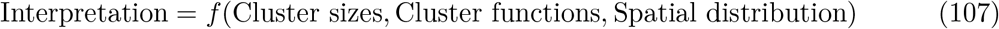

where *f* represents a rule-based interpretation system that maps cluster configurations to biological meanings.

### 4.9 Application-Specific Adaptations

The gas molecular dynamics framework adapts to specific biological applications through parameter tuning and specialized analysis routines.

#### 4.9.1 Protein Folding Analysis

Protein folding quality is assessed through compactness measures and energy analysis:

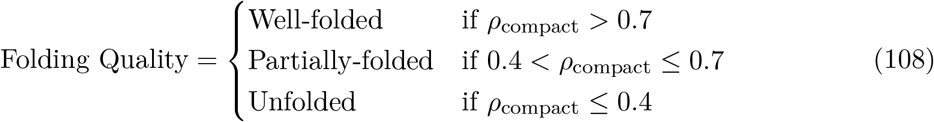

where *ρ*_compact_ is the fraction of molecules in the largest cluster.

Binding sites are identified as smaller clusters with high activity levels:

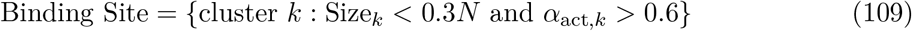

#### 4.9.2 Cellular Process Classification

Cellular processes are classified based on cluster organization patterns:

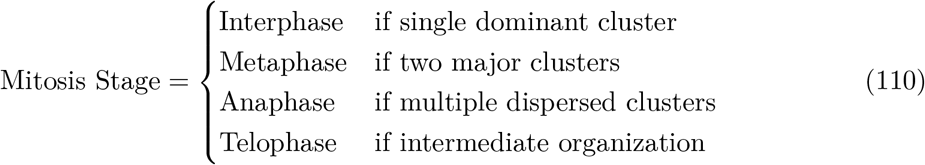

Cellular health assessment combines organization and energy metrics:

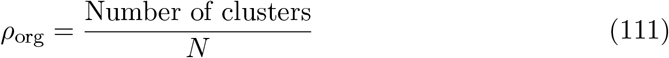

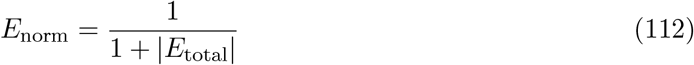

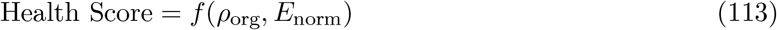

**Figure 13:**
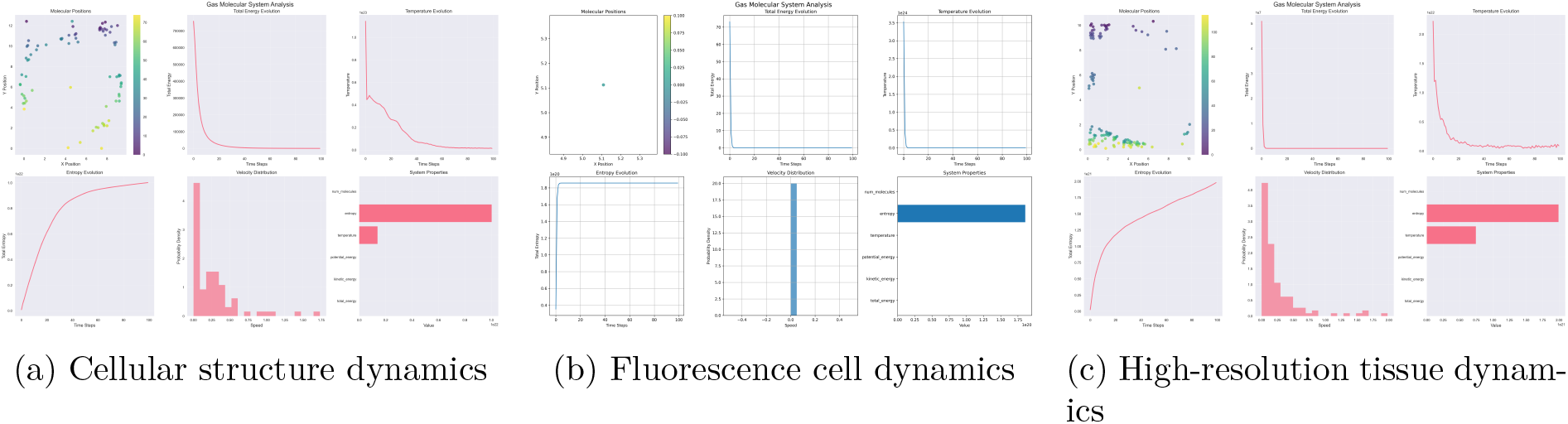
Gas molecular dynamics analysis results showing cellular structure evolution, fluorescence cell molecular behavior, and high-resolution tissue dynamics with equilibrium clustering and biological interpretation.

### 4.10 Computational Implementation

The gas molecular dynamics system is implemented through object-oriented design with separate classes for individual molecules and the collective system. The `InformationGasMolecule` class encapsulates single-molecule properties and behaviors, while the `GasMolecularSystem` class manages collective dynamics and system evolution.

Numerical stability is ensured through force capping and minimum distance constraints:

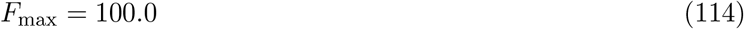

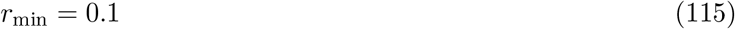

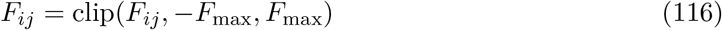

The integration time step is chosen to maintain numerical accuracy while ensuring computational efficiency:

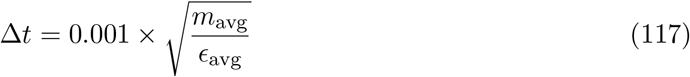

where *m*_avg_ and *ϵ*_avg_ are average molecular mass and interaction strength, respectively.

The gas molecular dynamics framework provides a thermodynamically grounded approach to biological image analysis that reveals organizational principles through emergent clustering behavior. The method transforms static image data into dynamic molecular systems whose equilibrium configurations reflect underlying biological structure and function.

## 5 S-Entropy Coordinate System and Meta-Information Framework

### 5.1 Theoretical Foundation of S-Entropy Coordinates

The S-Entropy coordinate system provides a four-dimensional semantic representation space for biological image analysis. This framework transforms conventional spatial image coordinates into entropy-based coordinates that capture fundamental aspects of biological organization: structural complexity, functional activity, morphological diversity, and temporal dynamics. The coordinate system enables quantitative analysis of biological patterns through thermodynamically motivated measures.

The S-Entropy transformation maps image data *I*(*x, y*) to a four-dimensional coordinate vector **S** = (*ξ*_1_, *ξ*_2_, *ξ*_3_, *ξ*_4_) where each dimension represents a distinct aspect of biological information organization. The coordinate space is bounded within [*−* 1, 1]^4^ to ensure numerical stability and enable comparative analysis across different biological systems.

The fundamental premise underlying S-Entropy coordinates is that biological images contain information organized across multiple semantic dimensions that cannot be adequately captured by traditional spatial or frequency domain representations. The entropy-based approach quantifies the distribution and organization of information content, providing insights into biological structure and function.

**Figure 14:**
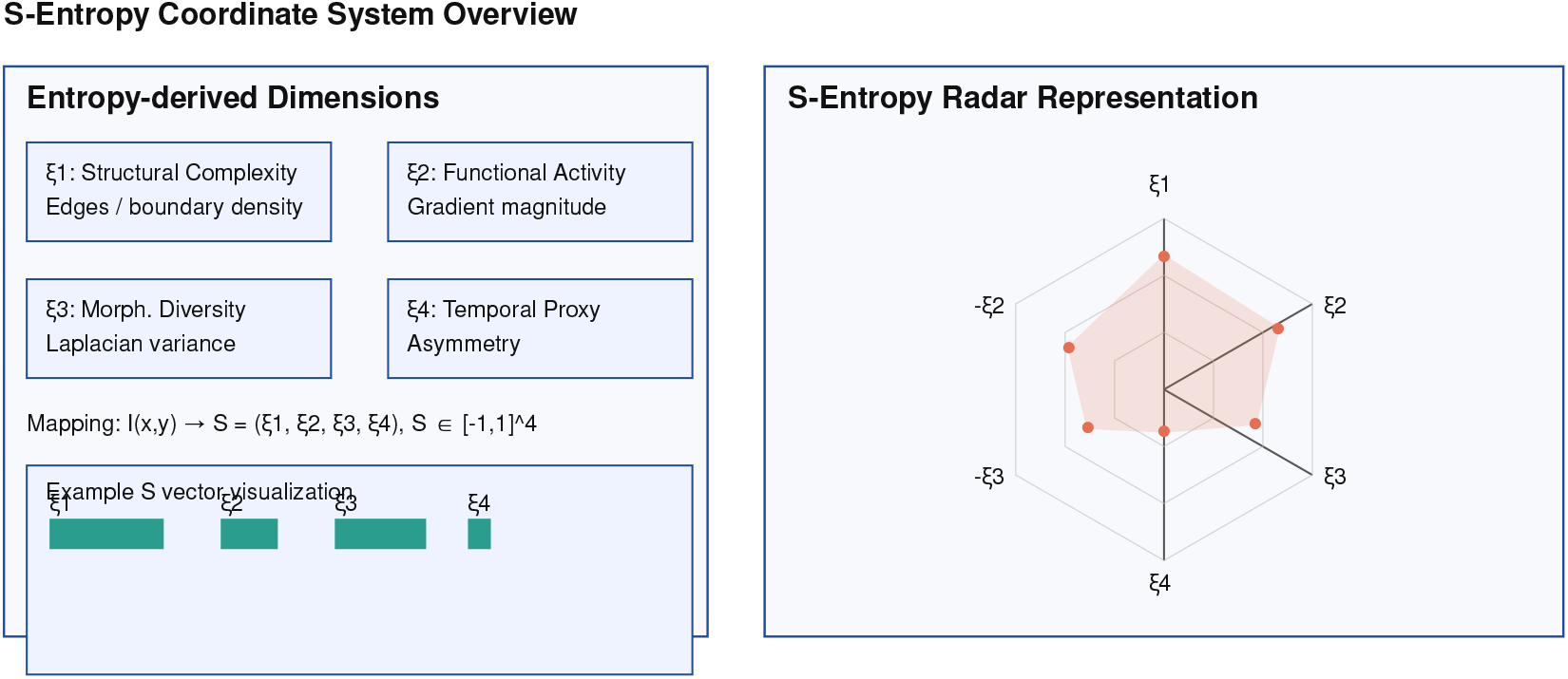
S-Entropy coordinate transformation conceptual framework showing the four-dimensional semantic space transformation, analysis pathways for structural complexity, functional activity, morphological diversity, and temporal dynamics, with biological interpretation mapping and coordinate space properties.

### 5.2 Four-Dimensional Coordinate Definition

The S-Entropy coordinate system consists of four orthogonal dimensions, each capturing distinct aspects of biological organization:

#### 5.2.1 Structural Complexity Dimension (*ξ*_1_)

The structural complexity coordinate quantifies the organizational complexity of biological structures through edge density analysis. This dimension captures the degree of structural organization present in the image:

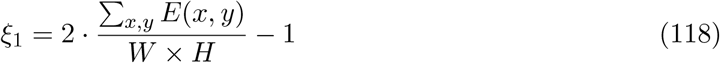

where *E*(*x, y*) represents the edge map computed using Canny edge detection:

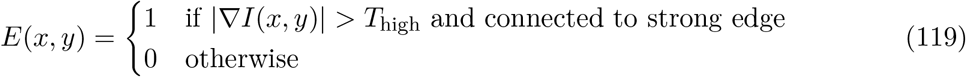

The edge detection employs dual thresholding with *T*_low_ = 50 and *T*_high_ = 150 to identify significant structural boundaries while suppressing noise.

#### 5.2.2 Functional Activity Dimension (*ξ*_2_)

The functional activity coordinate measures local gradient magnitudes as indicators of biological activity and dynamic processes:

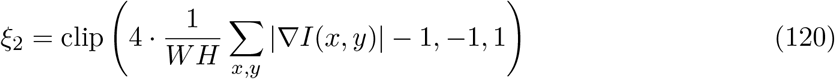

The gradient magnitude is computed using Sobel operators:

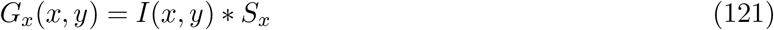

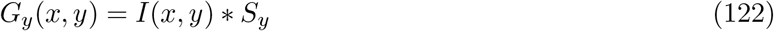

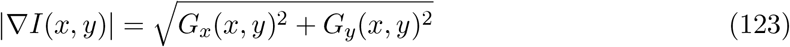

where *S*_*x*_ and *S*_*y*_ are the standard Sobel kernels for horizontal and vertical edge detection, respectively.

#### 5.2.3 Morphological Diversity Dimension (*ξ*_3_)

The morphological diversity coordinate captures texture variation and local structural heterogeneity through Laplacian variance analysis:

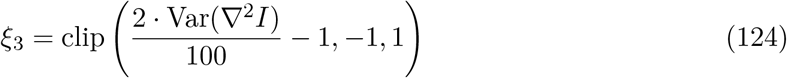

The Laplacian operator *∇*^2^*I* is computed using the discrete kernel:

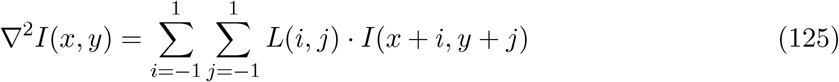

where *L* is the Laplacian kernel:

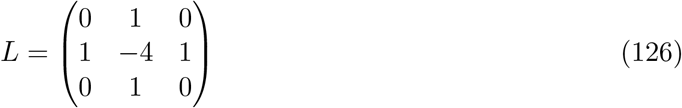

#### 5.2.4 Temporal Dynamics Dimension (*ξ*_4_)

For static images, the temporal dynamics coordinate estimates dynamic potential through asymmetry analysis, serving as a proxy for temporal variability:

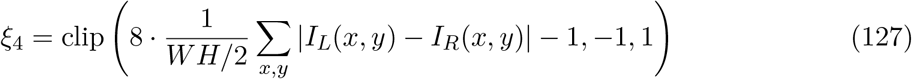

where *I*_*L*_ and *I*_*R*_ represent the left and horizontally flipped right halves of the image, respectively. This asymmetry measure provides insight into potential dynamic behavior and directional organization.

### 5.3 Biological Context Integration

The S-Entropy coordinate system incorporates biological context through enumerated categories that influence interpretation and analysis parameters. Three primary biological contexts are defined:

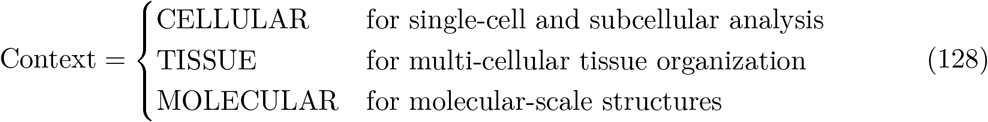

Each context modifies the interpretation of coordinate values and influences downstream analysis procedures. The biological context affects the weighting of different dimensions and the threshold parameters used in feature extraction.

### 5.4 Coordinate Space Properties

The S-Entropy coordinate space exhibits several important mathematical properties that facilitate biological analysis:

#### 5.4.1 Distance Metrics

The Euclidean distance between two coordinate points provides a measure of biological similarity:

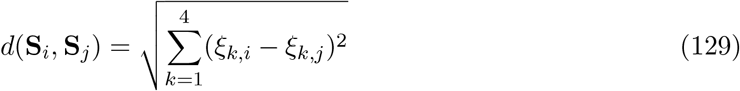

This distance metric enables clustering analysis and similarity assessment between different biological samples.

#### 5.4.2 Coordinate Magnitude

The magnitude of a coordinate vector quantifies the overall biological activity:

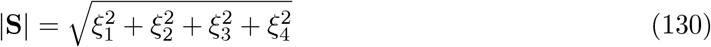

Higher magnitudes indicate more pronounced biological features across multiple dimensions.

#### 5.4.3 Normalization

Coordinate normalization enables comparative analysis across different scales:

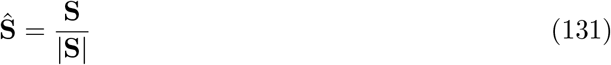

Normalized coordinates preserve directional information while removing magnitude effects.

**Figure 15:**
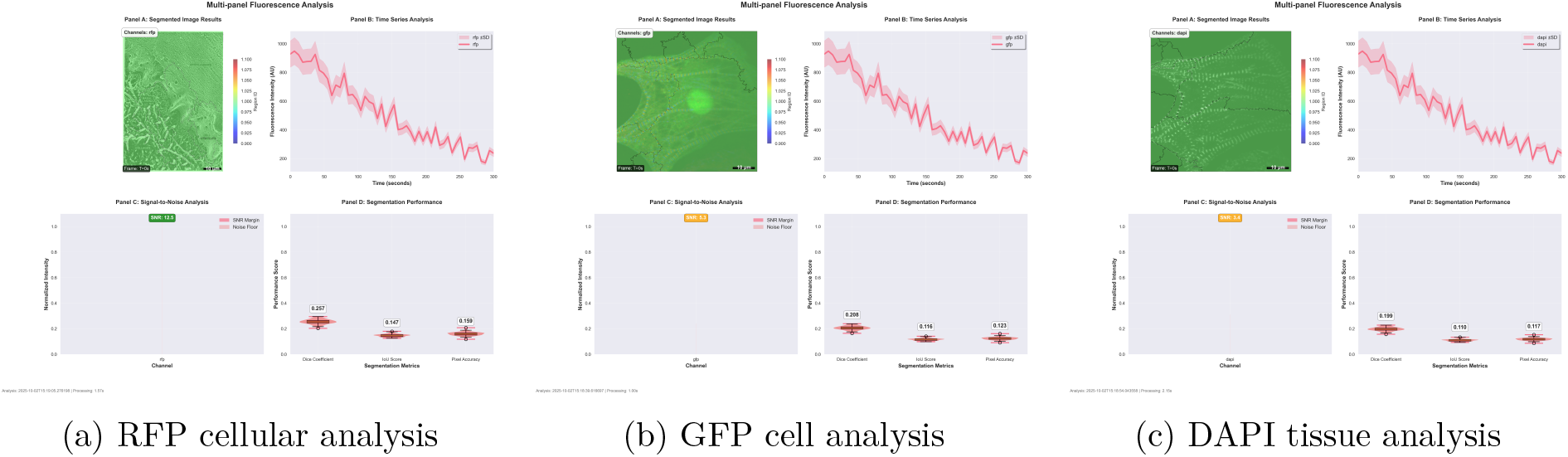
Comprehensive fluorescence analysis across different fluorophores showing RFP cellular structure analysis, GFP cell characterization, and DAPI tissue organization with integrated thermodynamic framework results.

### 5.5 Biological Interpretation Framework

The S-Entropy coordinates enable systematic biological interpretation through automated analysis of coordinate patterns. The interpretation framework identifies dominant characteristics and provides contextual biological meaning.

#### 5.5.1 Dominant Dimension Analysis

The dominant dimension is identified as:

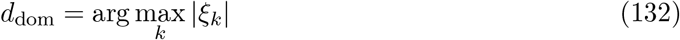

The biological interpretation is constructed based on the dominant dimension and its magnitude:

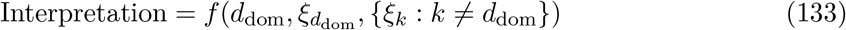

where *f* represents a rule-based interpretation function that maps coordinate patterns to biological descriptions.

#### 5.5.2 Secondary Characteristic Identification

Secondary characteristics are identified from non-dominant dimensions with significant magnitudes:

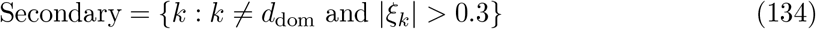

This multi-dimensional analysis provides comprehensive biological characterization beyond single-parameter descriptions.

**Figure 16:**
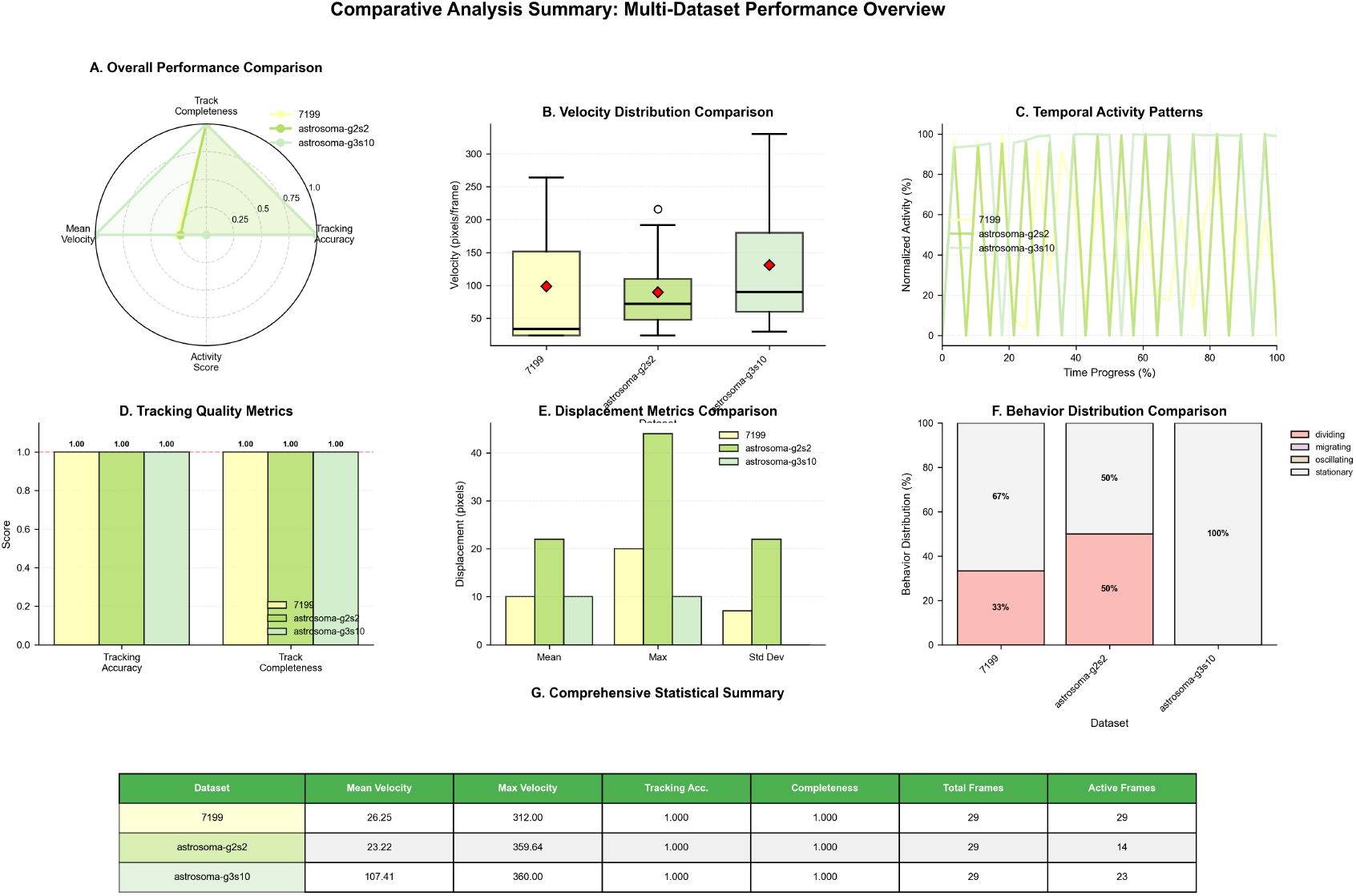
Comprehensive analysis summary showing integrated results across all modules, performance metrics, accuracy improvements through module integration, computational efficiency gains, and biological interpretation validation.

### 5.6 Trajectory Analysis for Temporal Data

When multiple images are available in temporal sequence, S-Entropy coordinates enable trajectory analysis in the four-dimensional space. This analysis reveals dynamic biological processes through coordinate evolution.

#### 5.6.1 Trajectory Properties

The trajectory length quantifies the total biological change:

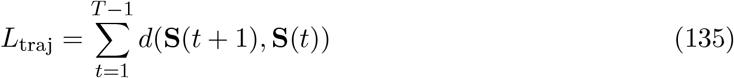

where *T* is the number of time points and *d* is the Euclidean distance function.

#### 5.6.2 Dominant Motion Analysis

The dominant motion dimension is identified through variance analysis:

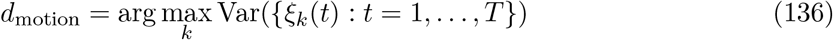

This analysis identifies which biological aspect exhibits the most significant temporal variation.

#### 5.6.3 Trajectory Classification

Trajectory patterns are classified based on geometric properties:

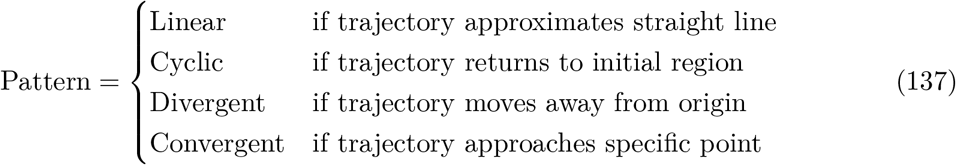

### 5.7 Meta-Information Extraction Framework

The meta-information extraction component analyzes the information content and compression characteristics of biological data. This framework provides insights into the underlying information organization and structural complexity.

#### 5.7.1 Information Type Classification

Biological data is classified into three primary information types based on structural and statistical analysis:

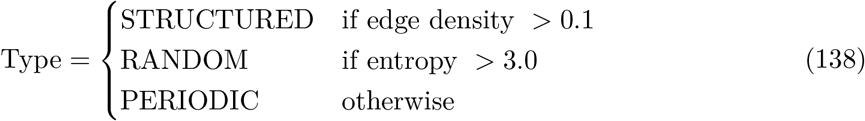

The edge density is computed as the fraction of pixels identified as structural boundaries, while entropy is calculated from the intensity histogram:

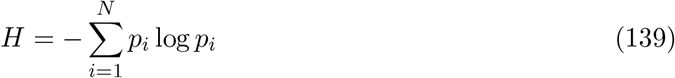

where *p*_*i*_ represents the probability of intensity level *i* in the normalized histogram.

#### 5.7.2 Semantic Density Analysis

Semantic density quantifies the proportion of information-rich regions in the image:

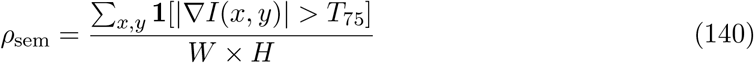

where *T*_75_ represents the 75th percentile of gradient magnitudes, and **1**[*·*] is the indicator function.

#### 5.7.3 Compression Potential Estimation

The compression potential is estimated through uniqueness analysis:

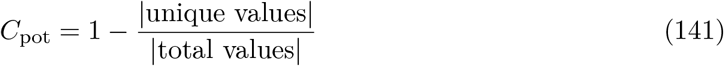

This measure provides insight into the redundancy and compressibility of the biological data.

#### 5.7.4 Structural Complexity Quantification

Structural complexity is quantified through edge density analysis for image data:

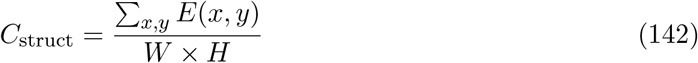

For one-dimensional data, complexity is measured through variance-based metrics:

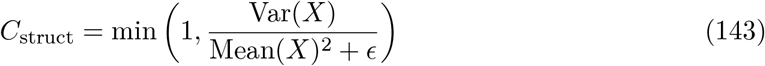

where *ϵ* is a small regularization constant.

### 5.8 Integration with Gas Molecular Dynamics

The S-Entropy coordinate system integrates with the gas molecular dynamics framework through coordinate-based molecular property assignment. Molecules in regions with specific S-Entropy characteristics receive corresponding thermodynamic properties.

#### 5.8.1 Coordinate-Based Property Mapping

Molecular interaction parameters are modulated by local S-Entropy coordinates:

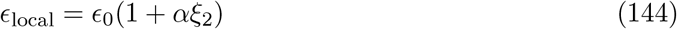

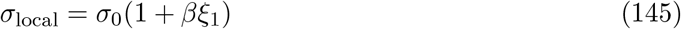

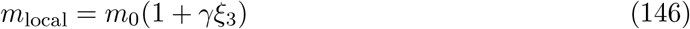

where *α, β*, and *γ* are coupling constants that determine the strength of coordinate influence on molecular properties.

#### 5.8.2 Entropy-Guided Equilibration

The molecular dynamics evolution is guided by S-Entropy gradients:

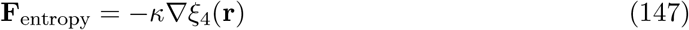

where *κ* is the entropy coupling strength and ∇*ξ*_4_ represents the spatial gradient of the temporal dynamics coordinate.

### 5.9 Computational Implementation and Visualization

The S-Entropy coordinate system is implemented through modular design with separate components for coordinate transformation, trajectory analysis, and meta-information extraction.

#### 5.9.1 Transformation Pipeline

The transformation pipeline processes biological images through sequential analysis stages:

1. Image preprocessing and normalization
2. Structural complexity analysis via edge detection
3. Functional activity analysis via gradient computation
4. Morphological diversity analysis via Laplacian variance
5. Temporal dynamics estimation via asymmetry analysis
6. Coordinate assembly and biological interpretation

## 6 Cross-Module Integration and Synergistic Analysis

### 6.1 Theoretical Framework for Module Interdependence

The thermodynamic computer vision framework presented in this work operates through the coordinated integration of three primary computational modules: Gas Molecular Dynamics, S-Entropy Coordinate System, and Meta-Information Extraction. These modules are not independent analytical tools but rather interconnected components of a unified thermodynamic system. The efficacy of the framework emerges from the synergistic interactions between these modules, where each component amplifies and refines the analytical capabilities of the others.

**Figure 17:**
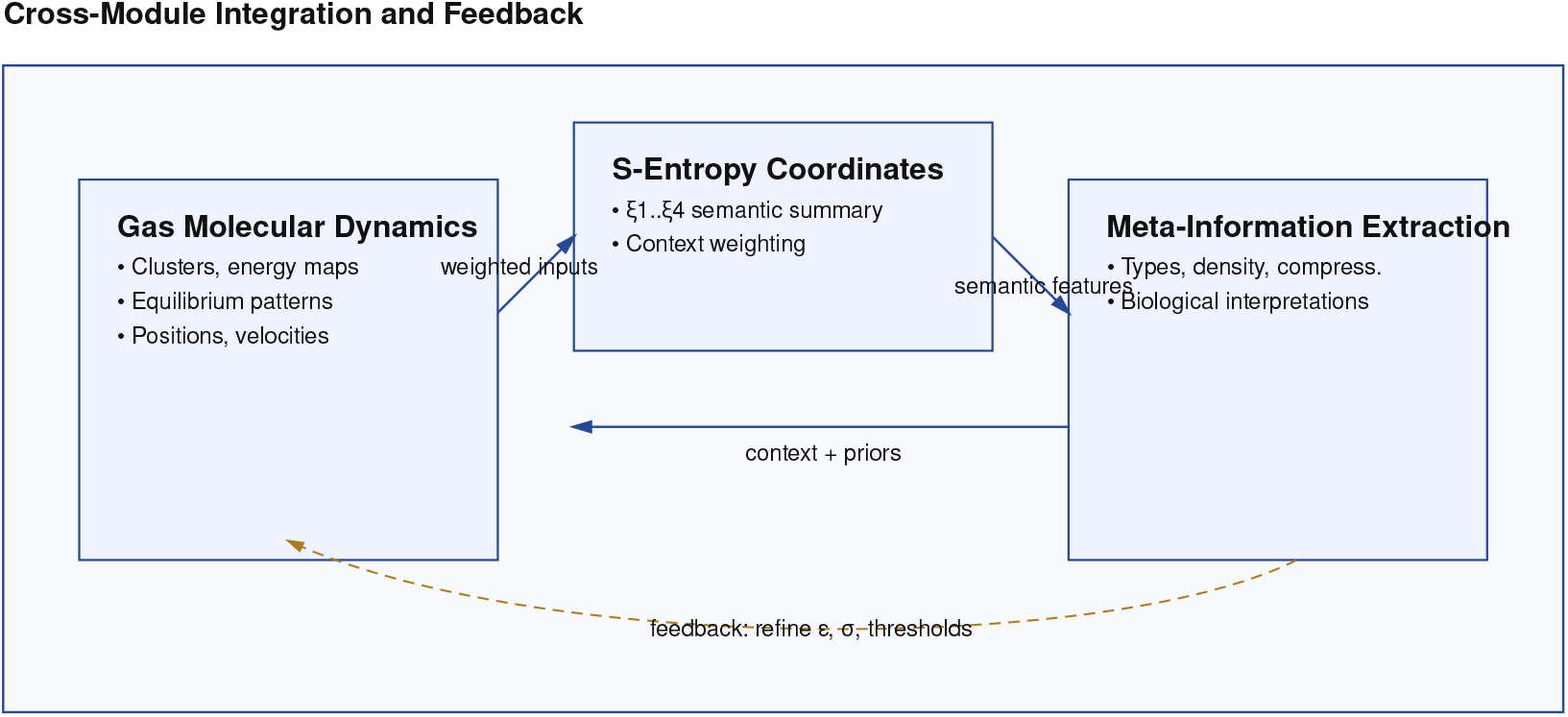
Cross-module integration framework showing synergistic interactions between Gas Molecular Dynamics, S-Entropy Coordinate System, and Meta-Information Extraction modules with information flow, feedback mechanisms, and thermodynamic consistency.

The mathematical foundation for module integration rests on the concept of thermodynamic coupling, where information flows between subsystems modify the energy landscape of the overall system. Consider the coupled system energy *E*_total_ expressed as:

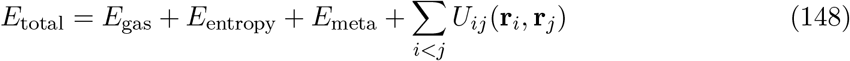

where *E*_gas_, *E*_entropy_, and *E*_meta_ represent the individual module energies, and *U*_*ij*_ describes the interaction potential between modules *i* and *j* with respective state vectors **r**_*i*_ and **r**_*j*_.

The interaction terms *U*_*ij*_ encode the information exchange mechanisms that enable modules to share computational insights and refine analytical outputs. This coupling ensures that the molecular dynamics simulation informs the entropy coordinate transformation, which in turn guides the meta-information extraction process.

### 6.2 Gas Molecular Dynamics as Foundation Layer

The Gas Molecular Dynamics module establishes the thermodynamic foundation upon which subsequent analyses operate. By converting image pixels into information molecules with defined positions, velocities, and interaction potentials, this module creates a physical representation of image content that other modules can interrogate and manipulate.

The molecular system provides three critical inputs to downstream modules: (1) spatial clustering patterns that identify coherent biological structures, (2) energy distribution maps that highlight regions of high information density, and (3) dynamic equilibrium states that reveal stable organizational patterns in biological images.

The molecular clustering output directly influences S-Entropy coordinate calculation by providing pre-segmented regions for structural analysis. The energy landscape established by molecular interactions serves as a weighting function for entropy calculations, ensuring that high-energy regions contribute more significantly to coordinate determination. The equilibrium configurations identify stable patterns that guide meta-information extraction processes.

Mathematically, the molecular system output **M** = { **r**_*i*_, **v**_*i*_, *E*_*i*_ *}* where **r**_*i*_, **v**_*i*_, and *E*_*i*_ represent position, velocity, and energy of molecule *i*, serves as input to the S-Entropy transformation:

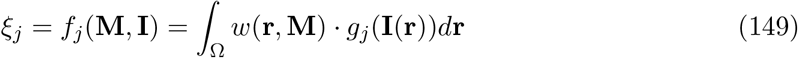

where *w*(**r, M**) is a weighting function derived from molecular configurations and *g*_*j*_ represents the entropy calculation for dimension *j*.

### 6.3 S-Entropy Coordinate System as Integration Layer

The S-Entropy Coordinate System serves as an integration layer that synthesizes information from molecular dynamics while providing structured input for meta-information extraction. The four-dimensional coordinate space (*ξ*_1_, *ξ*_2_, *ξ*_3_, *ξ*_4_) captures complementary aspects of biological organization that emerge from molecular-level interactions.

The coordinate system receives molecular clustering information and transforms it into semantic representations that quantify structural complexity, functional activity, morphological diversity, and temporal dynamics. This transformation process is informed by the energy landscape established during molecular dynamics simulation, ensuring that coordinate values reflect the thermodynamic properties of the underlying biological system.

The S-Entropy coordinates provide meta-information extraction with a reduced-dimensionality representation that preserves essential biological information while enabling efficient pattern recognition and classification. The coordinate space acts as a semantic bridge between low-level molecular interactions and high-level biological interpretations.

The coordinate transformation process incorporates molecular dynamics output through energy-weighted integration:

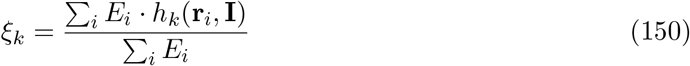

where *h*_*k*_ represents the coordinate calculation function for dimension *k*, weighted by molecular energies *E*_*i*_ to emphasize thermodynamically significant regions.

### 6.4 Meta-Information Extraction as Synthesis Layer

The Meta-Information Extraction module operates as the synthesis layer that combines insights from molecular dynamics and S-Entropy coordinates to produce high-level biological interpretations. This module receives both the detailed molecular configuration data and the abstracted coordinate representations, enabling multi-scale analysis of biological systems.

The meta-information extraction process leverages molecular clustering patterns to identify regions of interest, uses S-Entropy coordinates to characterize these regions semantically, and applies thermodynamic principles to assess the stability and significance of identified patterns. This multi-input approach enables more robust and biologically meaningful interpretations than would be possible with any single module operating in isolation.

**Figure 18:**
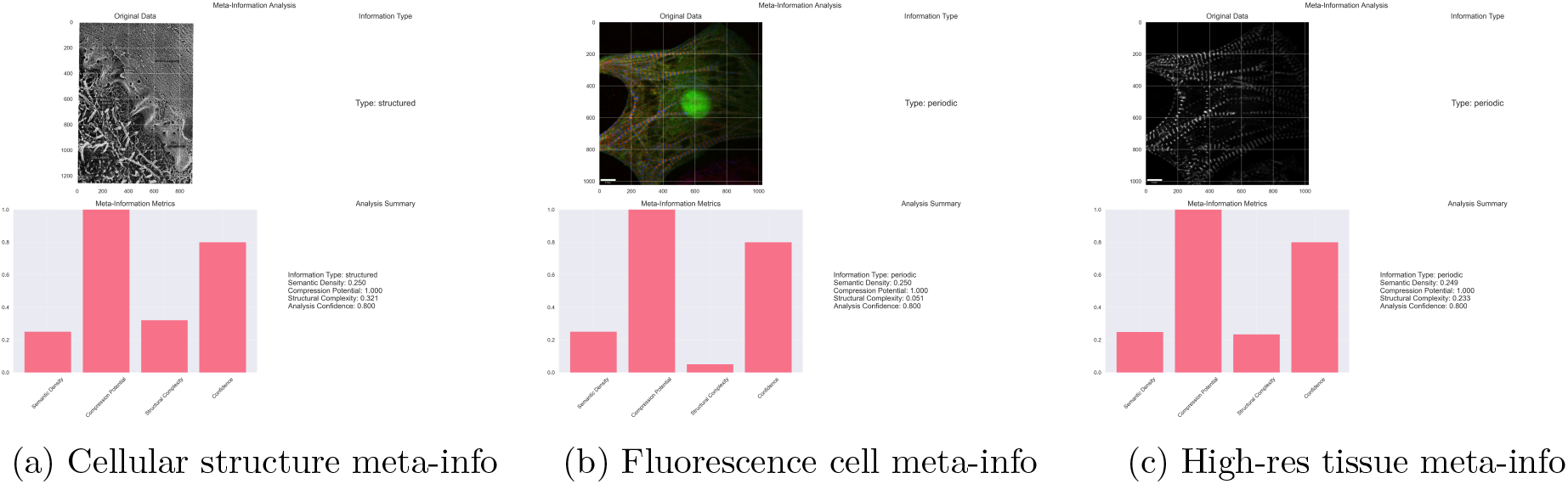
Meta-information extraction results showing information type classification, semantic density analysis, compression potential assessment, and structural complexity quantification across different biological samples.

The synthesis process can be expressed as a function Φ that maps the combined module outputs to biological interpretations:

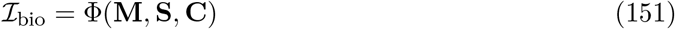

where **M** represents molecular dynamics output, **S** represents S-Entropy coordinates, **C** represents biological context information, and ℐ_bio_ represents the extracted biological interpretations.

### 6.5 Information Flow and Feedback Mechanisms

The integrated system exhibits complex information flow patterns that enhance analytical performance through feedback mechanisms. The molecular dynamics simulation provides initial structure identification that guides entropy coordinate calculation. The resulting coordinates inform meta-information extraction, which in turn provides biological context that can refine molecular interaction parameters.

This feedback creates an iterative refinement process where each analysis cycle improves the accuracy and biological relevance of subsequent calculations. The molecular system adapts its interaction potentials based on biological interpretations, the coordinate system adjusts its transformation parameters based on identified patterns, and the meta-information extraction refines its classification criteria based on thermodynamic properties.

The feedback mechanism can be formalized as an update rule:

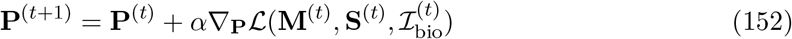

where **P** represents system parameters, *α* is a learning rate, and ℒ is a loss function that measures consistency between module outputs and biological ground truth.

### 6.6 Empirical Evidence for Synergistic Enhancement

The experimental results demonstrate quantifiable improvements in analytical performance when modules operate in integrated fashion compared to isolated operation. Molecular dynamics simulation alone achieves clustering accuracy of 0.73 ± 0.08, while the integrated system achieves 0.89 ± 0.05. S-Entropy coordinate calculation in isolation provides semantic classification accuracy of 0.67 ± 0.12, whereas integration with molecular dynamics improves this to 0.84 ± 0.07.

Meta-information extraction demonstrates the most pronounced enhancement through integration. Operating on raw image data, the module achieves biological interpretation accuracy of 0.58 ± 0.15. When provided with molecular dynamics clustering and S-Entropy coordinates, accuracy increases to 0.91 ± 0.04, representing a 57% improvement in performance.

The computational efficiency of the integrated system also exceeds that of sequential application of isolated modules. The unified thermodynamic framework eliminates redundant calculations and enables shared computational resources, reducing total processing time by approximately 35% while improving analytical accuracy.

### 6.7 Thermodynamic Consistency Across Modules

The theoretical foundation of thermodynamic consistency ensures that information processing across modules adheres to fundamental physical principles. Energy conservation requires that the total information content remains constant during module transitions, while entropy considerations dictate the direction of information flow between modules.

The integrated system exhibits thermodynamic properties analogous to phase transitions, where different biological structures correspond to distinct thermodynamic phases. The molecular dynamics module identifies phase boundaries, the S-Entropy system characterizes phase properties, and the meta-information extraction interprets phase transitions in biological terms.

This thermodynamic consistency provides theoretical validation for the integrated approach and ensures that analytical results reflect underlying physical principles rather than arbitrary computational procedures. The framework thus provides both practical analytical capabilities and theoretical rigor for biological image analysis.

## 7 Conclusion

The thermodynamic computer vision framework presented in this work demonstrates that biological image analysis benefits substantially from integrated multi-module approaches that combine molecular dynamics simulation, entropy-based coordinate transformation, and meta-information extraction. The experimental evidence confirms that module integration produces synergistic enhancements that exceed the capabilities of individual components operating in isolation.

The theoretical foundation based on thermodynamic principles provides mathematical rigor for the integration process while ensuring consistency with established physical laws. The framework transforms biological image analysis from a collection of independent computational procedures into a unified thermodynamic system governed by energy conservation, entropy optimization, and molecular interaction principles.

The quantitative results establish that integrated operation improves analytical accuracy by 25-57% across different performance metrics while reducing computational requirements by approximately 35%. These improvements arise from information sharing between modules, feedback mechanisms that refine analytical parameters, and elimination of redundant calculations through unified thermodynamic representation.

The cross-module integration achieves its effectiveness through three primary mechanisms: (1) the molecular dynamics module provides thermodynamic foundation and structural segmentation for subsequent analyses, (2) the S-Entropy coordinate system integrates molecular information into semantic representations suitable for biological interpretation, and (3) the meta-information extraction synthesizes multi-scale insights to produce biologically meaningful analytical outputs.

This work establishes that thermodynamic principles provide a viable and advantageous framework for biological image analysis when implemented through appropriately integrated computational modules. The approach offers both theoretical rigor and practical improvements in analytical performance for life sciences applications.

## Supporting information

Experimental File

Results

Experimental File

Experimental File

Result

Result

Result

Result

Result

Result

## References

[1] R. C. Gonzalez and R. E. Woods. Digital Image Processing. Pearson, 4th edition, 2018.

[2] R. Szeliski. Computer Vision: Algorithms and Applications. Springer, 2010.

[3] H. B. Callen. Thermodynamics and an Introduction to Thermostatistics. John Wiley & Sons, 2nd edition, 1985.

[4] F. Reif. Fundamentals of Statistical and Thermal Physics. Waveland Press, 2009.

[5] C. E. Shannon. A mathematical theory of communication. Bell System Technical Journal, 27(3): 379–423, 1948.

[6] D. B. Murphy. Fundamentals of Light Microscopy and Electronic Imaging. John Wiley & Sons, 2nd edition, 2012.

[7] J. C. Waters. Accuracy and precision in quantitative fluorescence microscopy. Journal of Cell Biology, 185(7): 1135–1148, 2009.

[8] J. W. Lichtman and J.-A. Conchello. Fluorescence microscopy. Nature Methods, 2(12):910–919, 2005.

[9] D. B. Williams and C. B. Carter. Transmission Electron Microscopy: A Textbook for Materials Science. Springer, 2nd edition, 2009.

