## Supplementary figures and images for "On the Thermodynamic Consequences of Oscillatory Dynamics in Image Processing: Application of Gas Molecule Models and Hierarchical Meta-Information Data Structures in Life Sciences Image Analysis"

### Experimental File

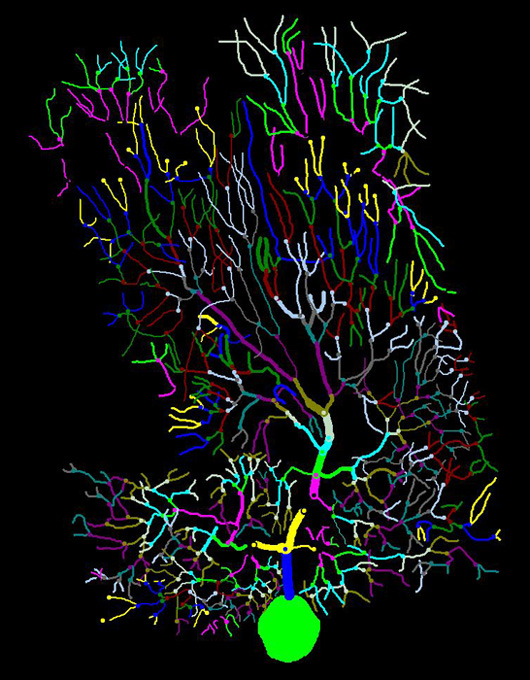

### Experimental File

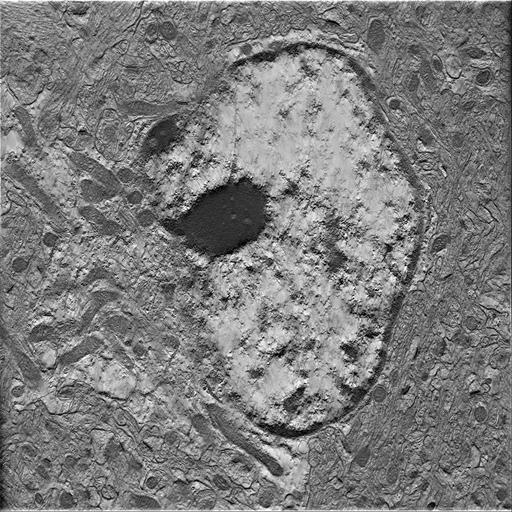

### Experimental File

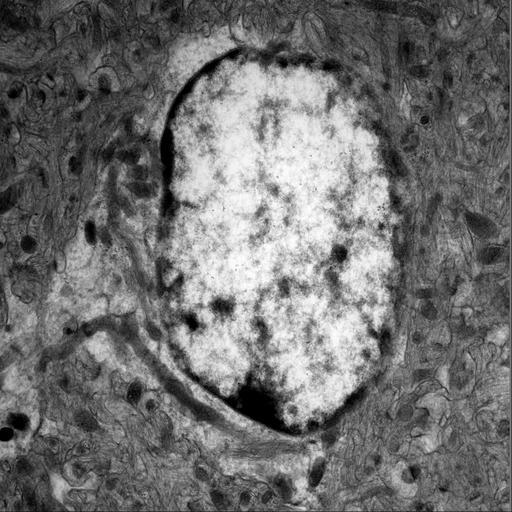
